# HIPPO signaling resolves embryonic cell fate conflicts during establishment of pluripotency *in vivo*

**DOI:** 10.1101/316539

**Authors:** Tristan Frum, Tayler Murphy, Amy Ralston

## Abstract

During mammalian development, the challenge for the embryo is to override intrinsic cellular plasticity to drive cells to distinct fates. Here, we unveil novel roles for the HIPPO signaling pathway segregates pluripotent and extraembryonic fates by controlling cell positioning as well as expression of *Sox2,* the first marker of pluripotency in the mouse early embryo. We show that maternal and zygotic YAP1 and WWTR1 repress *Sox2* while promoting expression of the trophectoderm gene *Cdx2* in parallel. Yet, *Sox2* is more sensitive than *Cdx2* to *Yap1/Wwtr1* dosage, leading cells to a state of conflicted cell fate when YAP1/WWTR1 activity is moderate. Remarkably, HIPPO signaling activity resolves conflicted cell fate by repositioning cells to the interior of the embryo, independent of its role in regulating *Sox2* expression. Rather, HIPPO antagonizes apical localization of Par complex components PARD6B and aPKC. Thus, negative feedback between HIPPO and Par complex components ensure robust lineage segregation.

## Introduction

During embryogenesis cells gradually differentiate, adopting distinct gene expression profiles and fates. In mammals, the first cellular differentiation is the segregation of trophectoderm and inner cell mass. The trophectoderm, which comprises the polarized outer surface of the blastocyst, will mainly produce cells of the placenta, while the inner cell mass will produce pluripotent cells, which are progenitors of both fetus and embryonic stem cells. Understanding how pluripotent inner cell mass cells are segregated from non-pluripotent cells therefore reveals how pluripotency is induced in a naturally occurring setting.

Progenitors of inner cell mass are first morphologically apparent at the 16-cell stage as unpolarized cells residing inside the morula (reviewed in Frum and Ralston, 2018). However, at this stage, pluripotency genes such as *Oct4* and *Nanog*, do not specifically label inside cells (Dietrich and Hiiragi, 2007; Niwa et al., 2005; Palmieri et al., 1994; Strumpf et al., 2005). Thus, the first cell fate decision has been studied mainly from the perspective of trophectoderm specification because the transcription factor CDX2, which is essential for trophectoderm development (Strumpf et al., 2005), is expressed specifically in outer cells of the 16-cell embryo (Ralston and Rossant, 2008), and has provided a way to distinguish future trophectoderm cells from non-trophectoderm cells. Knowledge of CDX2 as a marker of trophectoderm cell fate enabled the discovery of mechanisms that sense cellular differences in polarity and position in the embryo, and then respond by regulating expression of *Cdx2* (Nishioka et al., 2009). However, the exclusive study of *Cdx2* regulation does not provide direct knowledge of how pluripotency is established because the absence of *Cdx2* expression does not necessarily indicate acquisition of pluripotency. As such, our understanding of the first cell fate decision in the early mouse embryo is incomplete.

In contrast to other markers of pluripotency, *Sox2* is expressed specifically in inside cells at the 16-cell stage, and is therefore the first marker of pluripotency in the embryo (Guo et al., 2010; Wicklow et al., 2014). The discovery of how *Sox2* expression is regulated in the embryo provide therefore provide unique insight into how pluripotency is first established *in vivo*. Genes promoting expression of *Sox2* in the embryo have been described (Cui et al., 2016; Wallingford et al., 2017). However, it is currently unclear how expression of *Sox2* becomes restricted to inside cells. Interestingly, *Sox2* is restricted to inside cells by a *Cdx2*-independent mechanism (Wicklow et al., 2014), which differs from *Oct4* and *Nanog*, which are restricted to the inner cell mass by CDX2 (Niwa et al., 2005; Strumpf et al., 2005). Thus, *Sox2* and *Cdx2* are regulated in parallel, leading to complementary inside/outside expression patterns. However, it is not known whether *Sox2* is regulated by the same pathway that regulates *Cdx2* or whether a distinct pathway could be in use.

The expression of *Cdx2* is regulated by members of the HIPPO signaling pathway. In particular, the HIPPO pathway kinases LATS1/2 become active in unpolarized cells located deep inside the embryo, where they antagonize activity of the YAP1/WWTR1/TEAD4 transcriptional complex that is thought to promote expression of *Cdx2* (Anani et al., 2014; Cockburn et al., 2013; Hirate et al., 2013; Kono et al., 2014; Korotkevich et al., 2017; Leung and Zernicka-Goetz, 2013; Lorthongpanich et al., 2013; Mihajlović and Bruce, 2016; Nishioka et al., 2009, 2008; Posfai et al., 2017; Rayon et al., 2014; Watanabe et al., 2017; Yagi et al., 2007; Zhu et al., 2017). In this way, the initially ubiquitous expression of *Cdx2* becomes restricted to outer trophectoderm cells. However, the specific requirements for *Yap1* and *Wwtr1* in the regulation of *Cdx2* has been inferred from overexpression of wild type and dominant-negative variants, neither of which provide the standard of gene expression analysis that null alleles can provide. Nonetheless, the roles of *Yap1* and *Wwtr1* in regulating expression of *Sox2* have not been investigated. Here, we evaluate the roles of maternal and zygotic YAP1/WWTR1 in regulating expression of *Sox2* and cell fate during blastocyst formation.

## Results

### Patterning of *Sox2* is ROCK-dependent

To identify the mechanisms regulating *Sox2* expression during blastocyst formation, we focused on how *Sox2* expression is normally repressed in the trophectoderm to achieve inside cell-specific expression. We previously showed that SOX2 is specific to inside cells in the absence of the trophectoderm factor CDX2 (Wicklow et al., 2014), suggesting that mechanisms that repress *Sox2* in the trophectoderm act upstream of *Cdx2*. Rho-associated, coiled-coil containing protein kinases (ROCK1 and 2) are thought to act upstream of *Cdx2* because embryos developing in the presence of a ROCK-inhibitor (Y-27632, ROCKi) exhibit reduced *Cdx2* expression (Kono et al., 2014). Additionally, quantitative RT-PCR showed that *Sox2* mRNA levels are elevated in ROCKi-treated embryos (Kono et al., 2014), suggesting that ROCK1/2 activity leads to transcriptional repression of *Sox2*. However, the role of ROCK1/2 in regulating the spatial expression of *Sox2* has not been investigated.

To evaluate the roles of ROCK1/2 in patterning *Sox2* expression, we collected 8-cell stage embryos prior to embryo compaction (E2.5), and then cultured these either in control medium or in the presence of ROCKi for 24 hours (Fig. 1A). Embryos cultured in control medium exhibited normal cell polarity, evidenced by the apical localization of PARD6B and basolateral localization of E-cadherin (CDH1) in outside cells (Fig. 1B, C) as expected (Vestweber et al., 1987; Vinot et al., 2005). Additionally, SOX2 was detected only in inside cells in control embryos (Fig. 1C, D). By contrast, embryos cultured in ROCK inhibitor exhibited defects in cell polarity (Fig. 1B’, C’), consistent with prior studies (Kono et al., 2014). Interestingly, in ROCK inhibitor-treated embryos, we observed ectopic SOX2 expression in cells located on the outer surface of the embryo (Fig. 1C’, D), indicating that ROCK1/2 participates in the pathway responsible for repressing expression of *Sox2* in the trophectoderm.

**Figure 1.**
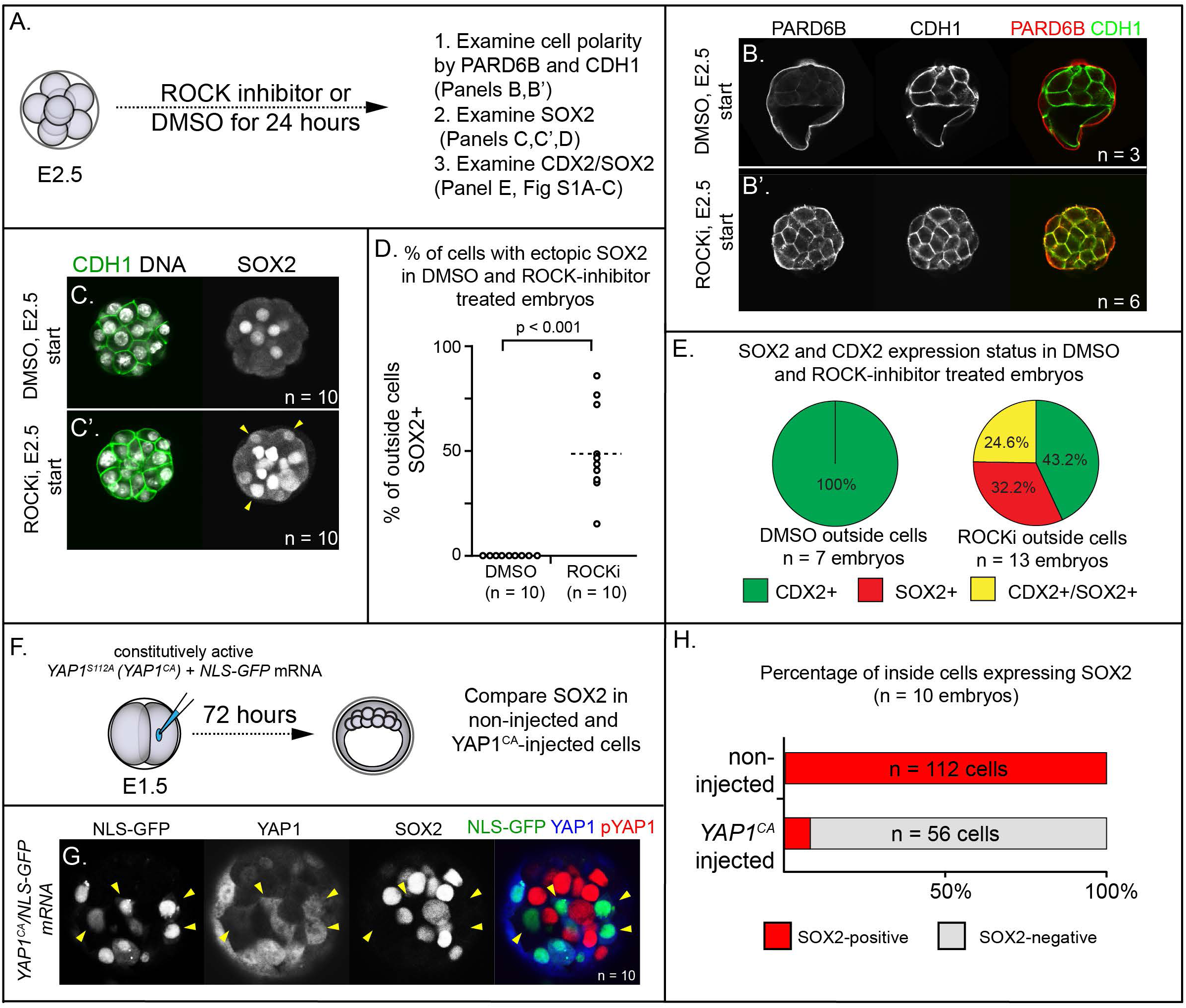
ROCK1/2 and nuclear YAP1 repress expression of SOX2. A) Experimental design: embryos were collected at E2.5 and treated with ROCK inhibitor Y-27632 (ROCKi) or DMSO (control) for 24 hours. B-B’) Confocal images of apical (PARD6B) and basolateral (CDH1) membrane components in control and ROCKi-treated embryos. As expected, PARD6B and CDH1 are mislocalized to the entire cell membrane of all cells in ROCKi-treated embryos, demonstrating effective ROCK inhibition (n = number of embryos examined). C-C’) In control embryos, SOX2 is detected only in inside cells, while in ROCKi-treated embryos, SOX2 is detected in inside and outside cells (arrowheads, outside cells; n = embryos). D) Quantification of ectopic SOX2 detected in outside cells of control and ROCKi-treated embryos (p, student’s t-test, n = embryos). E) SOX2 and CDX2 staining in outside cells of control and ROCKi-treated embryos. ROCK-inhibitor treatment leads to outside cells with mixed lineage marker expression (CDX2+/SOX2+). F) Experimental design: embryos were collected at E1.5 and one of two blastomeres injected with mRNAs encoding *YAP^CA^* and *GFP*. Embryos were cultured for 72 hours, fixed, and then analyzed by immunofluorescence and confocal microscopy. G) SOX2 is detected non-injected inside cells. SOX2 is not detected in *YAP^CA^-* overexpressing inside cells (arrowheads), n = embryos. H) Across multiple embryos, all non-injected inside cells express SOX2, whereas the vast majority of *YAP^CA^*-injected inside cells fail to express SOX2.

To scrutinize the identity of outside positioned SOX2-positive cells in ROCK-inhibited embryos, we co-stained an additional cohort of control and ROCKi-treated embryos with CDX2 and SOX2 and compared the overlap of lineage marker expression. In control embryos, CDX2 was detected only in outside cells (Fig. S1A) as expected at this stage (Ralston and Rossant, 2008; Strumpf et al., 2005). In ROCKi-treated embryos, CDX2 expression levels were reduced (Fig. S1A’) as was the proportion of outside cells in which CDX2 was detected (Fig. S1B), as previously reported (Kono et al., 2014).

However, among outside cells, a substantial proportion coexpressed CDX2 and SOX2 in ROCK-inhibited embryos compared with controls (Fig. 1E and S1A), suggesting that ROCK inhibition leads to an increase in outside cells of mixed lineage. Since SOX2 expression does not regulate expression of CDX2 (Wicklow et al., 2014), these observations suggest that ROCK1/2 activity regulate these genes through parallel mechanisms. We next sought to identify mediators that act downstream of ROCK1/2 to repress expression of *Sox2* in the trophectoderm.

### YAP1 is sufficient to repress expression of SOX2 in the inner cell mass

Several direct and indirect targets of ROCK1/2 kinases in the early embryo have been described (Alarcon and Marikawa, 2018; Shi et al., 2017). Among these is YAP1, a transcriptional partner of TEAD4 (Nishioka et al., 2009), since ROCK activity is required for the nuclear localization of YAP1 (Kono et al., 2014). Notably, *Tead4* is required to repress expression of *Sox2* in the trophectoderm (Wicklow et al., 2014), consistent with the possibility that YAP1 partners with TEAD4 to repress *Sox2* expression in the trophectoderm. To test this hypothesis, we overexpressed a constitutively active variant of YAP1 (YAP1^CA^). Substitution of alanine at serine 112 leads YAP1 to be constitutively nuclear and constitutively active (YAP1^CA^ hereafter) (Dong et al., 2007; Nishioka et al., 2009; Zhao et al., 2007). We injected mRNAs encoding *YAP1^CA^* and *GFP* into one of two blastomeres at the 2-cell stage, and then cultured these to the blastocyst stage (Fig. 1F). This mosaic approach to overexpression permitted comparison of *YAP1^CA^-* overexpressing with non-injected cells, which served as internal negative controls. We first examined localization of YAP1 in these embryos at the morula stage, with the expectation that YAP1 would be detected in nuclei of both inside and outside cells in *YAP1^CA^*-overexpressing cells (Nishioka et al., 2009). As expected, YAP1 was observed in nuclei of all *YAP1^CA^*-overexpressing cells (Fig. S1B, C). We next evaluated the consequences of ectopic nuclear YAP1 on expression of SOX2 in inside cells. We observed a conspicuous decrease in the proportion of *YAP1^CA^*-overexpressing inside cells lacking detectable SOX2 (Fig. 1G, H). Therefore, nuclear YAP1 is sufficient to repress *Sox2* expression in the inner cell mass, indicative of a likely role for YAP1 in repressing expression of *Sox2* in the trophectoderm downstream of ROCK1/2.

### LATS kinase is sufficient to induce inside cell positioning

To functionally test of the role of YAP1 in repressing expression of *Sox2*, we injected one of two blastomeres with mRNA encoding LATS2 kinase, which inactivates YAP1 and, presumably, the related protein WWTR1 by phosphorylation, causing their cytoplasmic retention (Nishioka et al., 2008). We then examined expression of SOX2 after culturing embryos to the blastocyst stage (Fig. 2A), predicting that LATS2 kinase would induce the ectopic expression of *Sox2* in outside cells. Surprisingly, we observed that almost all *Lats2*-overexpressing cells ended up within the inner cell mass by the blastocyst stage (Fig. 2B, C), in contrast to cells injected with GFP mRNA only, which contributed to both inner cell mass and trophectoderm. Notably, SOX2 was detected in all *Lats2*-overexpressing cells observed within the inner cell mass (Fig. 2D), suggesting that *Lats2*-overexpressing cells were not only localized to the inner cell mass but also position-appropriate regulation of *Sox2.*

**Figure 2.**
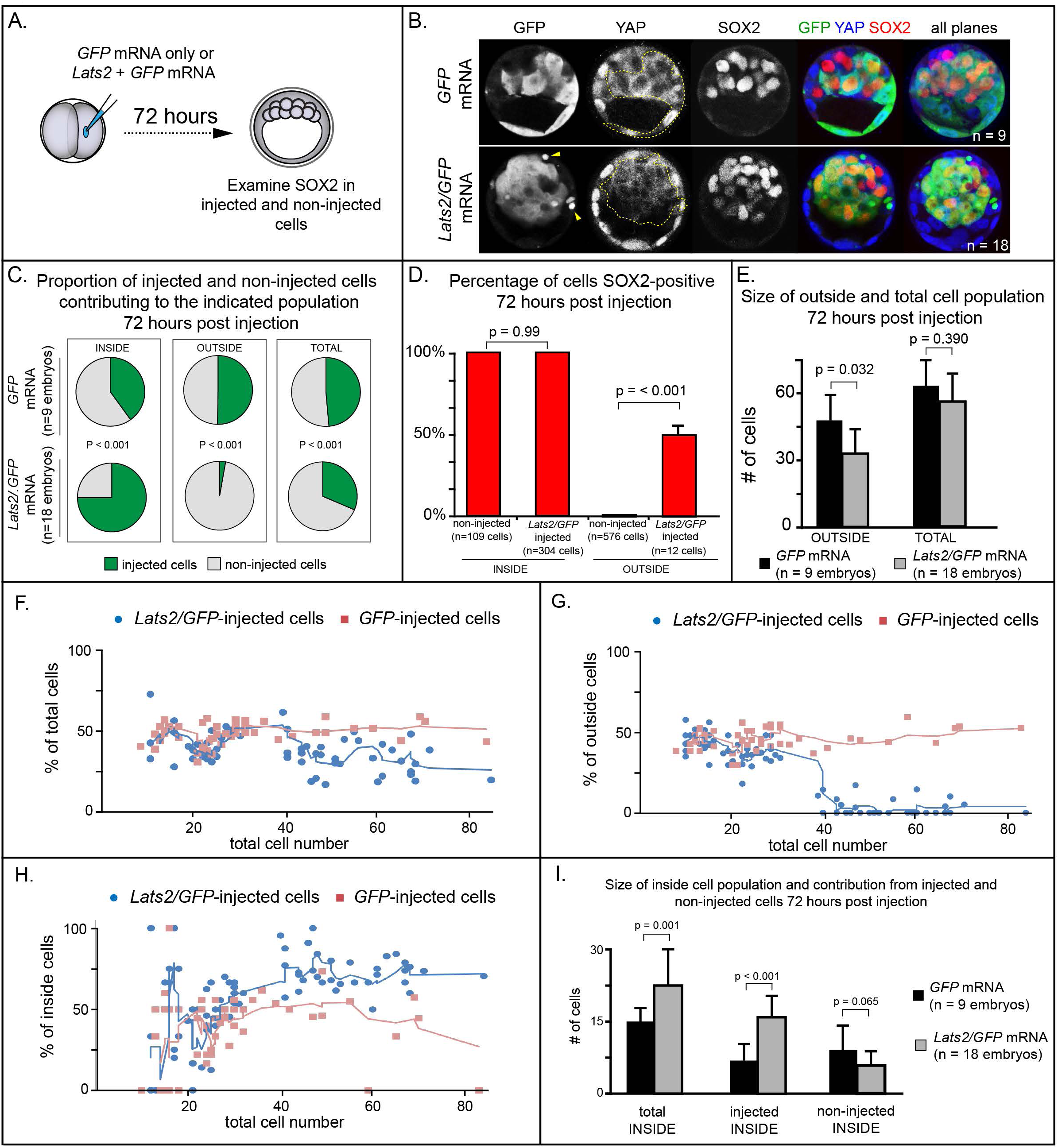
LATS2 kinase is sufficient to direct cells to inner cell mass fate. A) Embryos were collected at E1.5 and one of two blastomeres was injected with mRNAs encoding *Lats2* and *GFP* or *GFP* alone. Embryos were cultured for 72 hours, fixed, and then analyzed by immunofluorescence and confocal microscopy. B) Cells injected with *GFP* (dotted line) contributed to trophectoderm and inner cell mass, while cells injected with *Lats2* and *GFP* (dotted line) contributed almost exclusively to the inner cell mass, leaving only cellular fragments in the trophectoderm (arrows), suggestive of cell death (n = embryos). C) Proportion of inside, outside, and total cell populations across multiple embryos, which were comprised of non-injected cells, or cells injected with either *GFP* or *GFP/Lats2* mRNAs. Cells injected with *GFP/Lats2* were overrepresented within the inside cell population and underrepresented in the outside and total cell populations, relative to cells injected with *GFP* alone (P, chi-squared test). D) Percentage of SOX2-positive cells within non-injected and *GFP*-injected or *Lats2/GFP*-injected populations observed inside and outside of the embryo. SOX2 was detected in all of the Lats2/GFP-injected inside cells, and in half of the rare, *Lats2/GFP-* injected outside cells (same number of embryos as in panel C) (p, student’s t-test). E) Average number of outside and total cells per embryo. The average number of outside cells is reduced in embryos injected with *Lats2/GFP*, relative to GFP-injected (p, student’s t-test). F) Proportion of *GFP* and *Lats2/GFP*-injected cells, relative to total cell number, over the course of development to the ∼80-cell blastocyst (Solid lines = average of indicated data point and four previous data points). G) Data as shown in panel H, shown relative to outside cell number. H) Data as shown in panel H, shown relative to inside cell number. I) Contribution of injected and non-injected cells to the inside cell population, following injection with *GFP* or *Lats2/GFP.* Injection with *Lats2/GFP* increases the overall number of inside cells compared to injection with GFP only through increasing the number of injected cells contributing to the inside cell population, without affecting the number of non-injected cells contributing to the inside cell population (p, student’s t-test).

The strikingly increased prevalence of *Lats2*-overexpressing cells in the inner cell mass was also associated with a stark decrease in the number of *Lats2*-overexpressing cells detected within the trophectoderm and a decrease in the number of outside cells compared to embryos injected with *GFP* mRNA alone (Fig. 2C, E), suggesting that *Lats2*-overexpressing outside cells either internalize or undergo cell death. Furthermore, we observed cellular fragments within the trophectoderm of *Lats2*-overexpressing embryos (Fig. 2B, yellow arrowheads), as well as increased TUNEL staining in *Lats2-*overexpressing embryos compared to embryos injected with *GFP* mRNA only (Fig. S2A-B, D), consistent with increased death of *Lats2*-overexpressing cells.

In addition to detecting SOX2 in all *Lats2*-overexpressing cells located inside the embryo, SOX2 was also detected in rare *Lats2*-overexpressing cells that remained on the embryo surface (Fig. 2D). Therefore, LATS2 is sufficient to induce expression of SOX2 in cells regardless of their position within the embryo. Importantly, the kinase-dead variant of LATS2 (Nishioka et al., 2009), did not alter cell positioning, survival, or SOX2 expression (Fig. S3A, B), consistent with a previous report (Posfai et al., 2017). Thus, overexpressed LATS2 influences cell position and gene expression by modulating the activity of YAP1 and possibly WWTR1. We predicted that, if *Lats2* overexpression drove cells to adopt inner cell mass fate by influencing YAP1 and WWTR1 activity, then co-overexpression of *Yap1^CA^* would enable *Lats2*-overexpressing cells to contribute to trophectoderm. Consistent with this prediction, cooverexpression of *Lats2* and *Yap1^CA^* led to a significant decrease in the proportion of *Lats2-* overexpressing cells contributing to the inside cell position, and a concomitant increase in the proportion of *Lats2*-overexpressing cells remaining in the outside position (Fig. S3C-F). Moreover, cooverexpression of *Lats2* and *Yap1^CA^* reduced the number of TUNEL positive nuclei, consistent with *Yap1^CA^* rescuing survival of outside positioned *Lats2*-overexpressing cells (Fig. S2C-D). Collectively, these observations strongly suggest that LATS2 promotes inside cell positioning by regulating the activities of YAP1 and, likely, the related protein WWTR1.

To pinpoint when *Lats2*-overexpressing cells come to occupy the inside of the embryo, we performed a time course, examining the position of injected and non-injected cells from the 16-cell to the blastocyst stage (∼80 cells). Surprisingly, between the 16 and 32cell stages, the proportion of injected and non-injected cells in the total, outside, and inside cell populations were comparable whether embryos had been injected with *Lats2* and *GFP* or *GFP* mRNA alone (Fig. 2F-H). In embryos injected with *GFP* mRNA alone, the proportion of injected and non-injected cells making up the total, outside, and inside cell populations remained constant throughout the time course. In contrast, starting around the 32-cell stage, the average proportion of *Lats2*-overexpressing cells making up the inside population began to increase dramatically. This increase was associated with a decrease in the proportion *Lats2*-overexpressing cells making up the outside population, consistent with internalization of *Lats2*-overexpressing cells after the 32-cell stage (Fig 2G). After the 32-cell stage, *Lats2*-injected cells became underrepresented as a proportion of the total cell population (Fig. 2H), lending further support to the idea that *Lats2*-overexpressing cells that fail to internalize undergo cell death. Interestingly, the inside-skewed contribution of *Lats2*-overexpressing cells did not influence the ability of non-injected cells to contribute to the ICM (Fig. 2I), arguing that *Lats2*-overexpression drives inside positioning cell-autonomously. We therefore conclude that *Lats2* overexpression acts on cell position and survival around the time of blastocyst formation.

### LATS2 induces positional changes independent of *Sox2*

Our observation that *Lats2*-overexpression induces both the expression of SOX2 and cell repositioning to inner cell mass prompted us to investigate whether SOX2 itself drives cell repositioning downstream of *Lats2*. In support of this hypothesis, SOX2 activity has been proposed to bias inner cell mass fate (Goolam et al., 2016; White et al., 2016). We therefore investigated whether *Sox2* is required for the inner cell mass-inducing activity of LATS2 by overexpressing *Lats2* in embryos lacking maternal and zygotic *Sox2* (Fig. 3A), as previously described (Wicklow et al., 2014). However, we observed that *Lats2*-overexpressing cells were equally likely to occupy inside position in the presence and absence of *Sox2* (Fig. 3B, C). Furthermore, *Lats2*-overexpressing cells were equally unlikely to occupy outside position in the presence and absence of *Sox2* (Fig. 3D). Therefore, although *Lats2* overexpression is sufficient to induce expression of *Sox2*, LATS2 acts on cell positioning/survival independently of *Sox2*.

**Figure 3.**
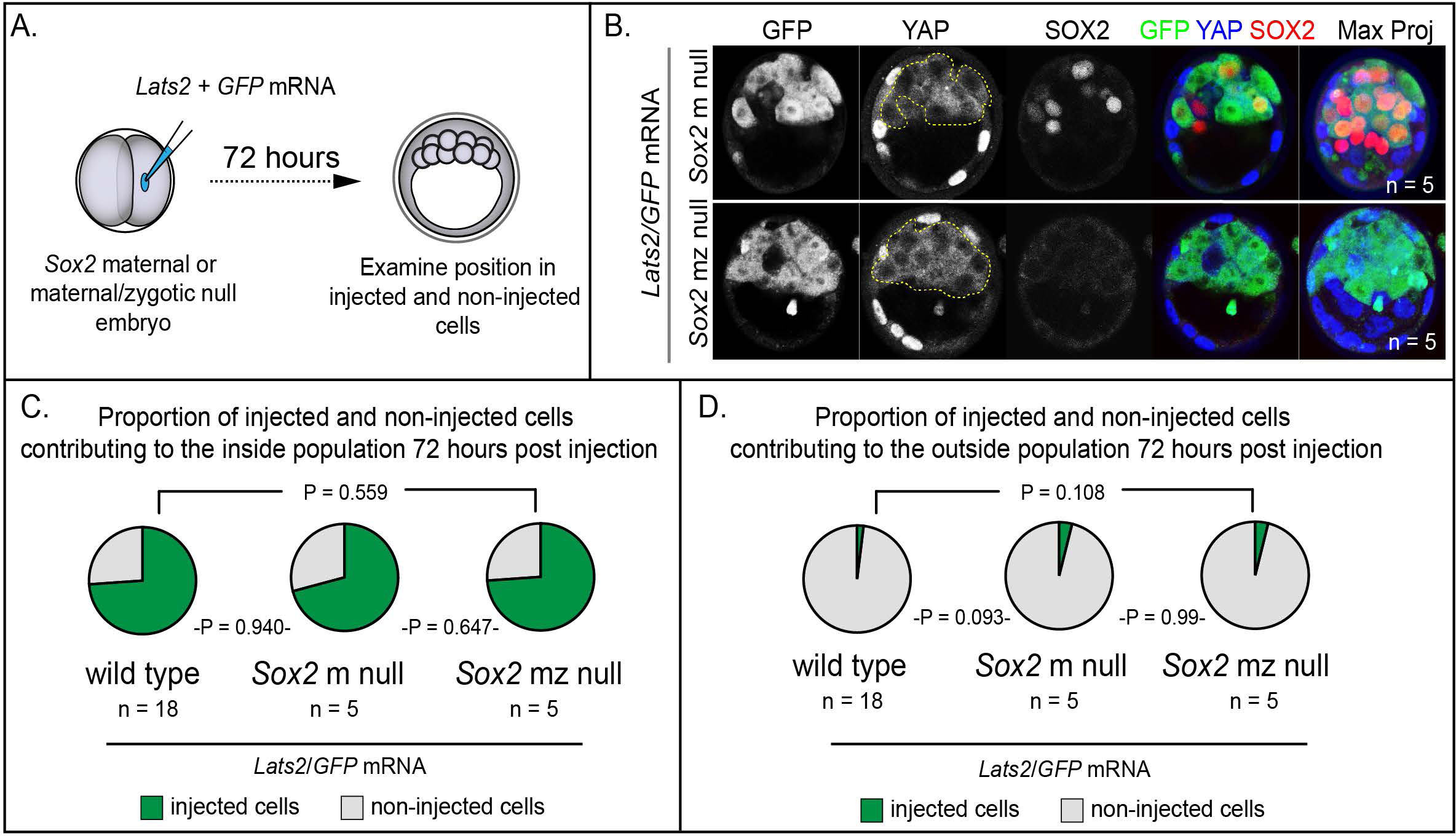
LATS2 directs inner cell mass fate independently of *Sox2*. A) *Lats2* and *GFP* or *GFP* alone were overexpressed in embryos lacking maternal or maternal and zygotic *Sox2.* B) *Lats2/GFP*-overexpressing cells (dotted line) contribute almost exclusively to the inner cell mass in the presence or absence of *Sox2* (n = embryos). C) Proportion of non-injected cells and cells injected with *Lats2/GFP* mRNAs contributing to inner cell mass in the indicated genetic backgrounds. No significant differences were observed based on embryo genotype, indicating that *Sox2* is dispensable for inside positioning by *Lats2*-overexpression (P, chi-squared test; n = embryos). D) Proportion of non-injected cells and cells injected with the indicated mRNAs contributing to trophectoderm in the indicated genetic backgrounds. No significant differences were observed based on embryo genotype (P, chi-squared test; n = embryos).

### LATS2 antagonizes formation of the apical domain

Trophectoderm cell fate has been proposed to be determined by apically localized membrane components that maintain the position of future trophectoderm cells on the embryo surface (Anani et al., 2014; Korotkevich et al., 2017; Maître et al., 2016, 2015; Samarage et al., 2015; Zenker et al., 2018). For example, the apical membrane components aPKC and PARD6B are required for maintaining outside cell position and trophectoderm fate (Alarcon, 2010; Dard et al., 2009; Hirate et al., 2015; Plusa et al., 2005). Because *Lats2* overexpression led cells to adopt an inside position, this raised the testable possibility that LATS2 antagonizes localization of aPKC and PARD6B.

Since *Lats2* overexpression leads to cell positioning starting around the 32-cell stage, we examined the localization of aPKCz and PARD6B in embryos just prior to the 32-cell stage. At this stage, apical membrane components PARD6B and aPKCz were detected at the apical membrane of non-injected outside cells and outside cells injected with *GFP* only (Fig. 4A-D). By contrast, most *Lats2*-overexpressing outside cells lacked detectable aPKCz and PARD6B (Fig. 4A-D). Therefore, LATS2 is sufficient to antagonize localization of key apical domain proteins in outside cells, providing a compelling mechanism for the observed repositioning of *Lats2*-overexpressing outside cells.

**Figure 4.**
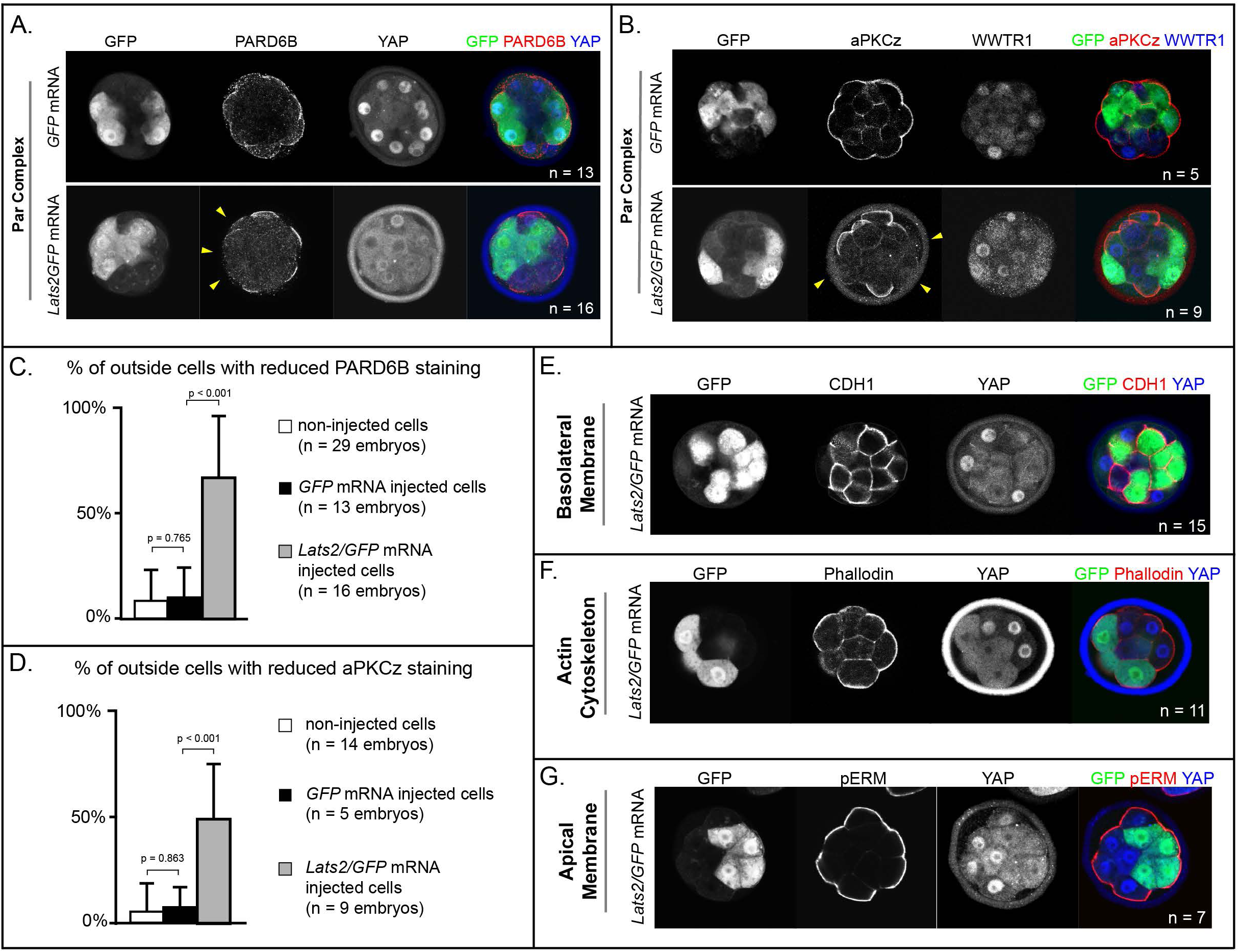
LATS2 antagonizes formation of the apical domain. A) In embryos at 16-32 cell stages, PARD6B is detectable in *GFP*-overexpressing and in non-injected cells, but not in *Lats2*-overexpressing cells (arrowheads, n = embryos). B) At 16-32 cell stages, aPKCz is detectable in *GFP*-overexpressing and in non-injected cells, but not in *Lats2*-overexpressing cells (arrowheads, n = embryos). C) Quantification of embryos shown in panel A (p, student’s t-test). D) Quantification of embryos shown in panel B (p, student’s t-test). E) At 16-32 cell stages, CDH1 is localized to the basolateral membrane in both *Lats2-* overexpressing and non-injected cells (n = embryos). F) At 16-32 cell stages, Phalloidin staining demonstrates that filamentous Actin is apically enriched in *Lats2*-overexpressing and non-injected cells (n = embryos). G) At 16-32 cell stages, pERM is localized to the apical membrane in both *Lats2-* overexpressing and non-injected cells (n = embryos).

We also examined other markers of apicobasal polarization in *Lats2*-overexpressing outside cells prior to the 32-cell stage. Curiously, other markers of apicobasal polarization were properly localized in all cells examined. For example, CDH1 was restricted to the basolateral membrane (Fig. 4E), while filamentous Actin and phospho-ERM were restricted to the apical domain in outside cells of both *Lats2*-overexpressing and non-injected outside cells (Fig. 4F, G). Thus, we propose that *Lats2*-overexpressing outside cells initially possess hallmarks of apicobasal polarization, but aPKC and PARD6B fail to properly localize, leading to their eventual depolarization and internalization.

### YAP1 and WWTR1 restrict *Sox2* expression to the inner cell mass

Our overexpression data suggested that the activities of YAP1 and WWTR1 are important for regulating cell fate and gene expression. Next, we aimed to test the requirement for *Yap1* and *Wwtr1* in embryogenesis. *Yap1* null embryos survive until E9.0 (Morin-Kensicki et al., 2006), suggesting that oocyte-expressed (maternal) *Yap1* (Yu et al., 2016), or the *Yap1* paralogue *Wwtr1* (Varelas et al., 2010) are important for preimplantation development. However, embryos lacking maternal and zygotic *Wwtr1* and *Yap1* have not been analyzed.

To generate embryos lacking maternal and zygotic *Wwtr1* and *Yap1,* we deleted *Wwtr1* and *Yap1* from the female germ line using mice carrying conditional alleles of *Wwtr1* and *Yap1* (Xin et al., 2013, 2011) and the female germ line-specific *Zp3Cre* (de Vries et al., 2000). We then crossed these females to males heterozygous for deleted alleles of *Wwtr1* and *Yap1* (see Methods). From these crosses, we obtained embryos lacking maternally provided *Wwtr1* and *Yap1* and either heterozygous or null for *Wwtr1* and/or *Yap1* (Table S1). At E3.25 (≤32 cells), SOX2 and CDX2 are normally mutually exclusive (Fig. 5A). However, with decreasing number of wild type zygotic alleles of *Wwtr1* and *Yap1*, we observed worsening phenotypes (Fig. 5B-F). In the complete absence of *Wwtr1* and *Yap1*, we observed a severe loss of CDX2 and expansion of SOX2 in outside cells (Fig. 5D-F), phenocopying *Lats2* overexpression. However, in embryos of intermediate genotypes, we observed expanded SOX2 and persistent, yet lower, expression levels of CDX2 (Fig. 5C, E-F). Thus, regulation of *Sox2* expression is more sensitive to *Wwtr1* and *Yap1* dosage than is *Cdx2*. Moreover, these observations indicate that intermediate doses of *Wwtr1* and *Yap1* produce outside cells expressing markers of mixed cell lineage at E3.25.

**Figure 5.**
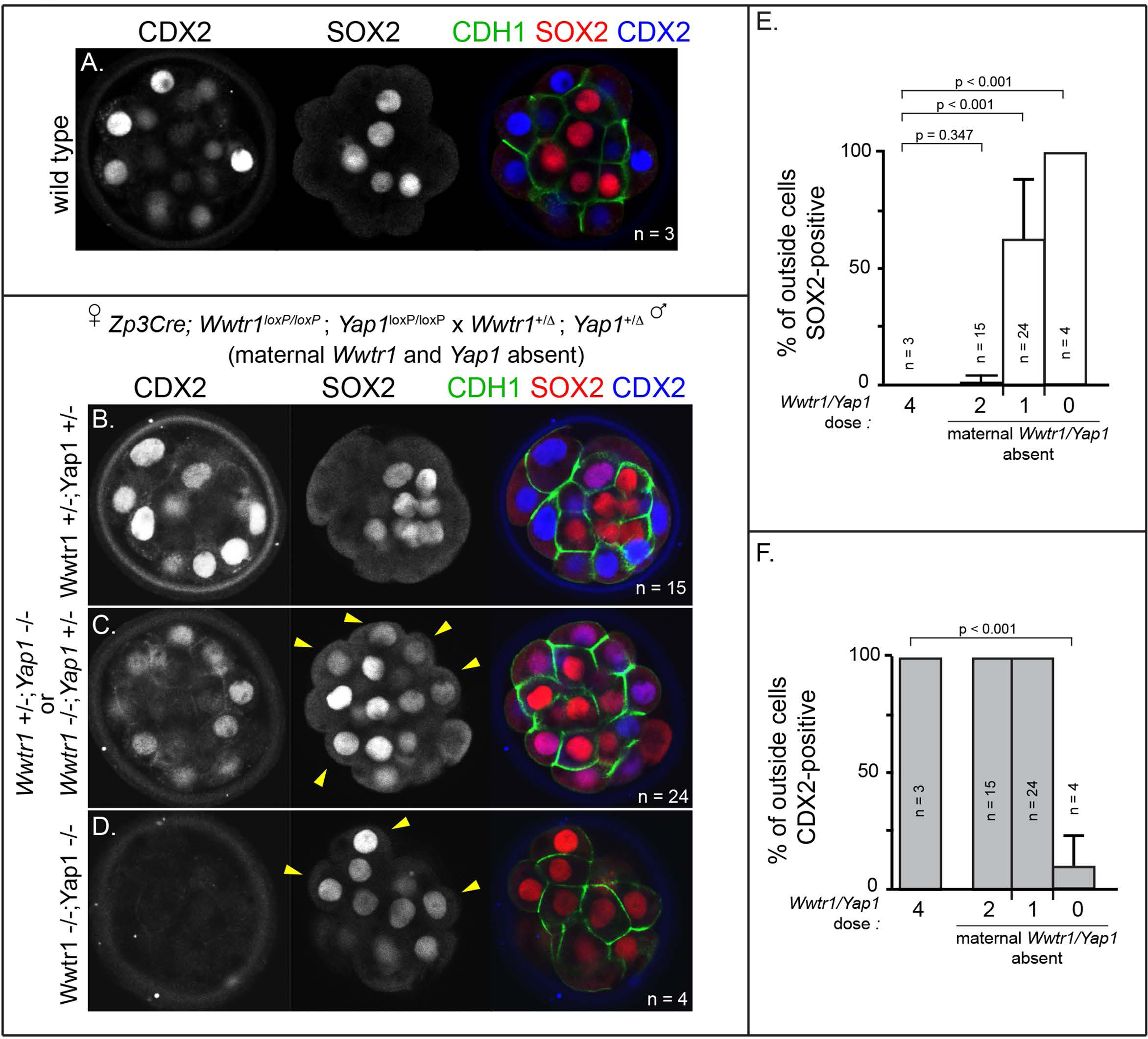
*Wwtr1* and *Yap1 are* required to repress SOX2 expression in outside cells. A) CDX2 and SOX2 in wild type embryos at E3.25 (16-32 cell stages). CDX2 staining is more intense in outside cells than inside cells and SOX2 staining is specific to inside cells (n = embryos). B) Embryos lacking maternal *Wwtr1* and *Yap1* with and heterozygous for *Wwtr1* and *Yap1* (which we consider to have 2 doses of WWTR1/YAP1) exhibit normal CDX2 and SOX2 expression (n = embryos). C) Embryos lacking maternal *Wwtr1* and *Yap1* and heterozygous for either *Wwtr1* or *Yap1* (1 dose of WWTR1/YAP1) exhibit a high degree of ectopic SOX2 in outside cells (arrowheads), but continue to express CDX2, although the levels appear reduced (n = embryos). D) Embryos lacking maternal and zygotic *Wwtr1* and *Yap1* (0 doses of WWTR1/YAP1) have the most severe phenotype, with a high degree of ectopic SOX2 in outside cells (arrowheads) and little or no detectable CDX2 (n = embryos). E) Quantification of the percentage of outside cells in which ectopic SOX2 is detected in the presence of decreasing dose of *Wwtr1* and *Yap1* (t = student’s t-test, n = embryos). F) Quantification of the percentage of outside cells in which CDX2 is detected in the presence of decreasing dose of *Wwtr1* and *Yap1* (t = student’s t-test, n = embryos).

### YAP1 and WWWTR1 maintain outside cell positioning

Based on our observations of *Lats2*-overexpressing embryos, we anticipated that defects in cell positioning in embryos lacking maternal and zygotic *Wwtr1* and *Yap1* could arise after the 32-cell stage. We therefore examined embryos lacking *Wwtr1* and *Yap1* at E3.75, when embryos possess more than 32-cells. Indeed, we observed skewed lineage contributions, correlating with the dosage of *Wwtr1* and *Yap1* (Fig. 6A-D). Embryos with one or fewer wild type alleles of *Wwtr1* or *Yap1* exhibited an increase in the number of inside cells and a reduction in the number of outside cells (Fig 6A-B), consistent with altered cell positioning.

**Figure 6.**
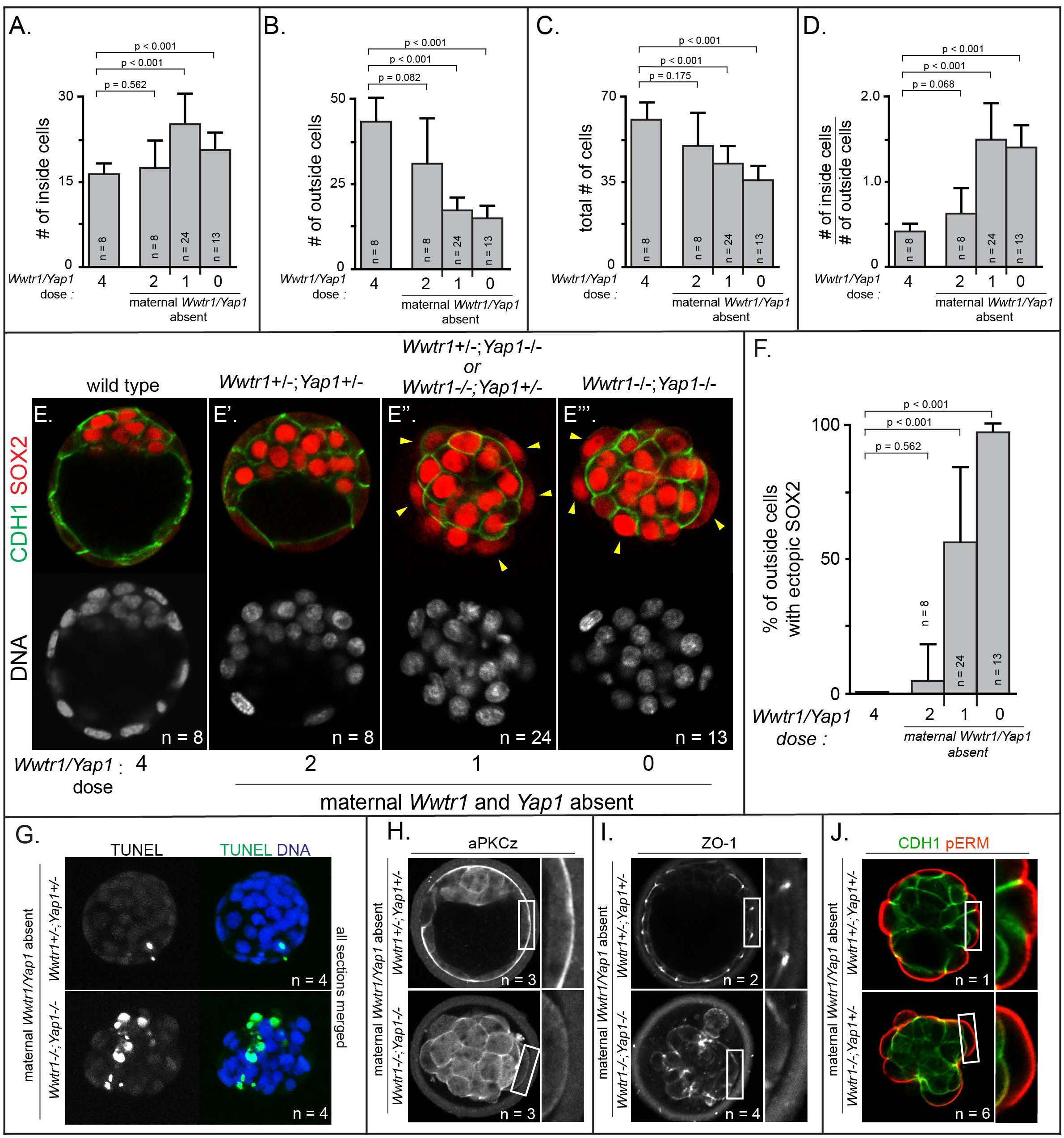
Positioning and epithelialization defects in embryos with *Wwtr1* and *Yap1* null alleles. A) Quantification of the average number of inside cells per embryo with decreasing dose of *Wwtr1* and *Yap1*. The number of inside cells increases as the dose of wild type *Wwtr1* and *Yap1* alleles is reduced (p, student’s t-test, n = embryos). B) Quantification of the average number of outside cells per embryo with decreasing dose of *Wwtr1* and *Yap1*. The number of outside cells decreases as the dose of wild type Wwtr1 and Yap1 alleles is reduced (p, student’s t-test, n = embryos). C) Quantification of the average number of total cells per embryos with decreasing dose of wild type zygotic *Wwtr1* and *Yap1*.The number of total cells decreases as the dose of wild type *Wwtr1* and *Yap1* is reduced (p, student’s t-test, n = embryos). D) Quantification of the average ratio of inside to outside cells per embryo with decreasing dose of *Wwtr1* and *Yap1*. The ratio of inside to outside cells increases as the dose of wild type *Wwtr1* and *Yap1* is reduced (p, student’s t-test, n = embryos). E) Wild type embryos at E3.75 exhibit inner cell mass-specific expression of SOX2 (n = embryos). E’) E3.75 embryos lacking maternal *Wwtr1* and *Yap1* and heterozygous for zygotic *Wwtr1* and *Yap1* cavitate and repress *Sox2* in outside cells, leading to inner cell mass-specific expression of SOX2 similar to wild type embryos (n = embryos). E’’) Embryos lacking maternal *Wwtr1* and *Yap1* but with only one wild type allele of *Wwtr1* or *Yap1* fail to cavitate and repress *Sox2* in outside cells, leading to ectopic SOX2 in outside cells (arrowheads, n = embryos). E’’’) Embryos lacking maternal and zygotic *Wwtr1* and *Yap1* fail to cavitate and repress *Sox2* in outside cells, leading to ectopic SOX2 in outside cells (arrowheads, n = embryos). F) Quantification of ectopic SOX2 detected in embryos such as those shown in panels E-E’’’. The percentage of outside cells with ectopic SOX2 increases as the dose of wild type *Wwtr1* and *Yap1* alleles is reduced (p, student’s t-test, n = embryos). G) TUNEL analysis of embryos lacking maternal *Wwtr1* and *Yap1* heterozygous for zygotic *Wwtr1* and *Yap1* or lacking maternal and zygotic *Wwtr1* and *Yap1.* Extensive TUNEL staining is observed in embryos lacking maternal and zygotic *Wwtr1* and *Yap1* indicative of cell death. Max projections of all confocal sections from a single embryo are shown (n = embryos). H) aPKCz staining in embryos lacking maternal *Wwtr1* and *Yap1,* either heterozygous for zygotic *Wwtr1* and *Yap1* or with no zygotic *Wwtr1* and *Yap1.* aPKC is not localized to the apical membrane of embryos with no zygotic *Wwtr1* and *Yap1* (n = embryos). I) ZO-1 staining in embryos lacking maternal *Wwtr1* and *Yap1,* either heterozygous for zygotic *Wwtr1* and *Yap1* or with no zygotic *Wwtr1* and *Yap1.* ZO-1 is disorganized in embryos with no zygotic *Wwtr1* and *Yap1,* suggesting that formation of a mature epithelium depends on *Wwtr1* and *Yap1* (n = embryos). J) pERM and CDH1 staining in embryos lacking maternal *Wwtr1* and *Yap1,* either heterozygous for zygotic *Wwtr1* and *Yap1* or with no zygotic *Wwtr1* and *Yap1.* pERM is localized to apical membranes and CDH1 to basolateral membranes regardless of the dose of wild type *Wwtr1* and *Yap1* alleles (n = embryos).

Although the average total number of cells was also reduced in these embryos (Fig. 6C), the reduction in total cell number did not alone account for the loss of cells on the outside of the embryo (Table S2). This observation suggested that, similar to *Lats2-* overexpressing cells, cells with reduced *Wwtr1* and *Yap1* exhibit an increased frequency of outside cell death, in addition to increased outside cell internalization. Consistent with this, embryos with one or fewer wild type alleles of *Wwtr1* or *Yap1* exhibited an increase in the ratio of inside to outside cells (Fig. 6D) and an increase in cells undergoing apoptosis by TUNEL assay (Fig. 6G and S6A, B).

Critically, the fewer outside cells apparent in embryos lacking *Wwtr1* and *Yap1*, which appeared stretched over the mass of inside cells, exhibited ectopic expression of SOX2 (Fig. 6E-F). Therefore, *Wwtr1/Yap1* repress inner cell mass fate, downstream of LATS kinases. Intriguingly, our data also indicate that WWTR1 is a more potent repressor of *Sox2* at E3.75 than YAP1 since embryos with a single wild type allele of *Wwtr1* had significantly fewer cells expressing ectopic SOX2 then embryos with a single wild type allele of *Yap1* (Fig. S4).

Since loss of *Wwtr1* and *Yap1* phenocopied *Lats2* overexpression in terms of *Sox2* expression, cell death, and cell repositioning, we next evaluated apical domain and cell polarization in outside cells of embryos lacking *Wwtr1* and *Yap1* at E3.75. We observed greatly reduced aPKC at the apical membrane of outside cells in embryos with one or fewer doses of *Wwtr1* or *Yap1* (Fig. 6H and S6C). In addition, we evaluated the localization of the tight junction protein ZO-1, which suggested failure in tight junction formation in embryos with 1 or fewer doses of *Wwtr1* and *Yap1* (Fig. 6I and S6D). Notably, however, other markers of apicobasal polarity, such as CDH1 and pERM were correctly localized in outside cells of mutant embryos at this stage (Fig. 6J and S6E). Our observations indicate that WWTR1 and YAP1 play a crucial role in the formation of the apical domain and maintaining the positioning and survival of outside cells while repressing expression of *Sox2*.

## Discussion

During preimplantation development, lineage-specific transcription factors are commonly expressed in ‘noisy’ domains before refining to a lineage-appropriate pattern (Simon et al., 2018). For example, *Oct4* and *Nanog* are expressed in both inner cell mass and trophectoderm until after blastocyst formation (Dietrich and Hiiragi, 2007; Strumpf et al., 2005). Similarly, CDX2 is detected in inner cell mass, as well as trophectoderm, until blastocyst stages (McDole and Zheng, 2012; Ralston and Rossant, 2008; Strumpf et al., 2005). In striking contrast to these genes, SOX2 is never detected in outside cells (Wicklow et al., 2014), indicating that robust mechanisms must exist to minimize noise and prevent its aberrant expression in trophectoderm. Here, we identify YAP1/WWTR1 as key components that repress *Sox2* expression in outside cells of the embryo. Notably, manipulations known to antagonize YAP1/WWTR1 activity, including chemical inhibition of ROCK and overexpression of LATS2 lead to ectopic expression of SOX2 in outside cells, reinforcing the notion that YAP1/WWTR1 activity are crucial for repression of *Sox2* in outside cells.

Additionally, we find that *Sox2* expression is more sensitive than is *Cdx2* to YAP1/WWTR1 activity, since intermediate doses of active YAP1/WWTR1 yields cells that coexpress both SOX2 and CDX2 (Fig. 7A). This observation is consistent with the fact that CDX2 is initially detected in inside cells of the embryo during blastocyst formation (Dietrich and Hiiragi, 2007; McDole and Zheng, 2012; Ralston and Rossant, 2008), where SOX2 is also expressed (Wicklow et al., 2014). Thus, inside cells could initially possess intermediate doses of active YAP1/WWTR1 at this early stage. By contrast, outside cells would have greatly reduced YAP1/WWTR1 activity, owing to elevated LATS activity. In this way, the HIPPO pathway ensures robust developmental transitions, by rapidly nudging SOX2-expressing cells into their correct and final positions inside the embryo (Fig. 7B).

**Figure 7.**
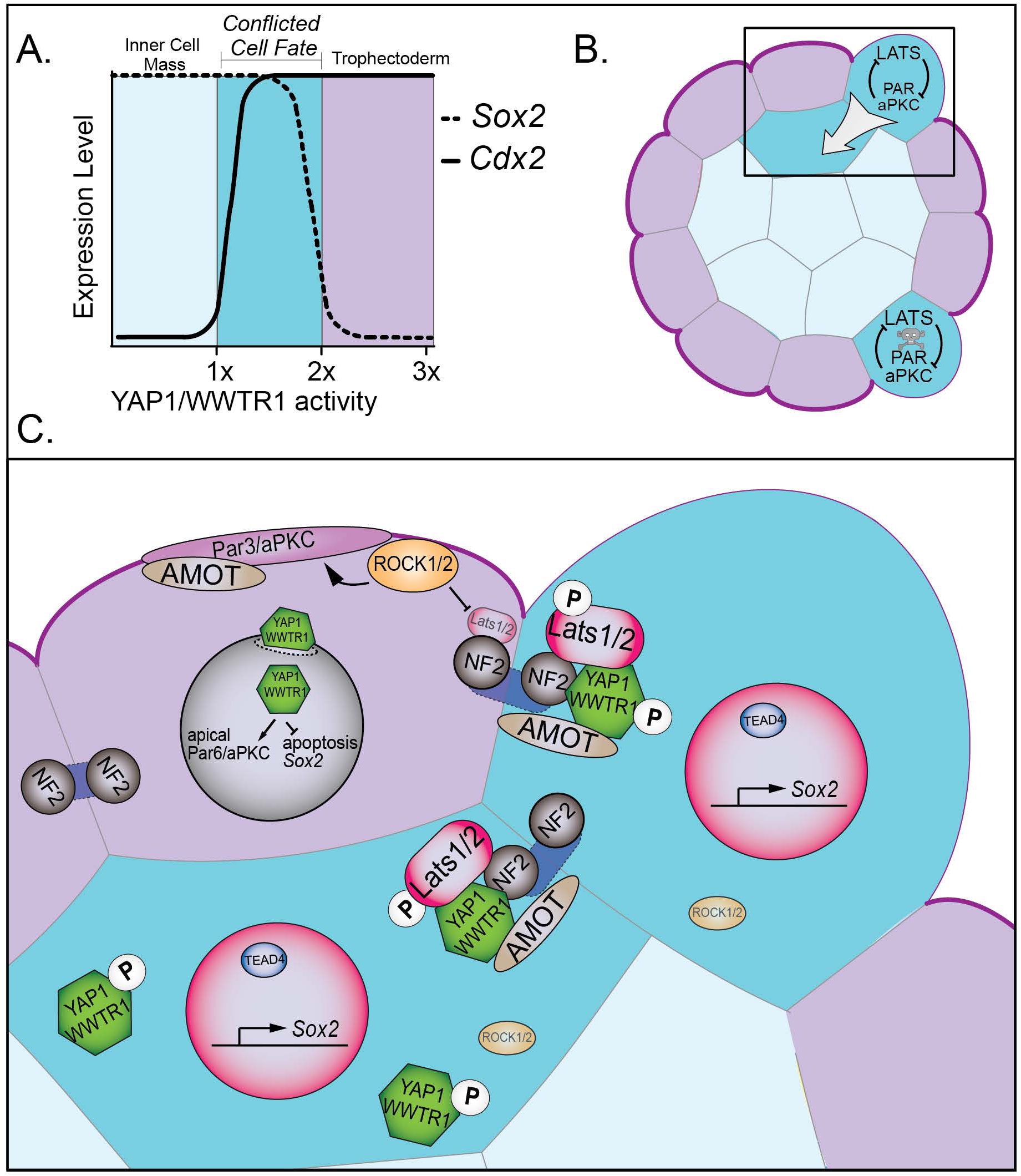
Resolution of cell fate conflicts in the preimplantation mouse embryo. A) The expression of *Sox2* and *Cdx2* is differentially sensitive to YAP1/WWTR1 activity, leading to co-expression of both lineage markers in cells when YAP1/WWTR1 activity levels are intermediate. B) During division from the 16- to the 32-cell stage, cells that inherit the apical membrane repress HIPPO signaling and maintain an outside position. However, cells that inherit a smaller portion of the apical membrane would initially elevate their HIPPO signaling. We propose that elevated HIPPO then feeds back onto polarity by further antagonizing PAR-aPKC complex formation, leading to a snowball effect on repression of *Sox2* expression, and thus ensuring that SOX2 is never detected in outside cells because these cells are rapidly internalized or apoptosed. C) A closeup of the boxed region in panel B. In most outside cells, low LATS2 activity enables high levels of YAP1/WWTR1 activity, which repress *Sox2* and apoptosis and promote *Cdx2* expression and apical localization of aPKC and PARD6B, which in turn repress the HIPPO pathway. In rare outside cells, LATS2 activity becomes elevated, leading to lower activity of YAP1/WWTR1, which then leads these cells to become internalized or to undergo apoptosis.

Consistent with our proposed model, the timing of HIPPO-induced cell internalization coincides with loss of cell fate plasticity around the 32-cell stage (Posfai et al., 2017). This timing also coincides with the formation of mature tight-junctions among outside cells (Sheth et al., 1997), which reinforce and intensify differences in HIPPO signaling activity between inside and outside compartments of the embryo (Hirate and Sasaki, 2014; Leung and Zernicka-Goetz, 2013). Our observations indicate that HIPPO signaling can, in turn, interfere with trophectoderm epithelialization. Therefore, we propose that HIPPO engages in a negative feedback loop with cell polarity components (Fig. 7B).

We propose two mechanisms by which HIPPO signaling eliminates cells from the trophectoderm, both of which are downstream of YAP1/WWTR1 (Fig. 7C). First, a small proportion of conflicted cells undergo cell death. This is in line with the observed increase in the level of apoptosis detected after the 32-cell stage (Copp, 1978). We showed that cell lethality due to elevated HIPPO can be rescued by increasing levels of nuclear YAP1, suggesting that YAP1 activity normally provides a pro-survival signal to trophectoderm cells, consistent with the proposed role of YAP1 in promoting proliferation in non-eutherian mammals (Frankenberg, 2018). Moreover, deletion of *Sox2* did not rescue survival of outside cells in which HIPPO signaling was artificially elevated, arguing that HIPPO resolves cell fate conflicts independently of lineage-specific genes.

The second way that conflicted cells are eliminated from the trophectoderm is that cells with elevated HIPPO signaling drive their own internalization. This is consistent with the observation that cells in which *Tead4* has been knocked down become internalized (Mihajlović et al., 2015). However, in contrast to *Tead4* loss of function, which preserves the polarization of outside cells (Mihajlović et al., 2015; Nishioka et al., 2008), we observed that *Yap1/Wwtr1* loss of function leads loss of apical PARD6D/aPKC. These observations suggest that YAP1/WWTR1 could partner with proteins other than TEAD4 to regulate apical domain formation. Consistent with this proposal, TEAD1 has been proposed to play an essential role in the early embryo (Sasaki, 2017). Nevertheless, since PARD6B/aPKC are essential for outside cell positioning (Dard et al., 2009; Hirate et al., 2015; Plusa et al., 2005), the loss of the apical domain could affect cell positioning in several ways. For instance, loss of PARD6B/aPKC would eventually lead to cell depolarization (Alarcon, 2010), which could influence any of the processes normally governing the formation of inside cells, such as oriented cleavage, cell contractility, or apical constriction (Korotkevich et al., 2017; Maître et al., 2016; Samarage et al., 2015). Identifying the downstream mechanisms by which HIPPO drives cells to inner cell mass will be a stimulating topic of future study.

Our studies also revealed that SOX2 does not play a role in cell positioning. This observation sheds light on a recent study, which showed that SOX2 dwells longer in select nuclei of four-cell stage embryos that are destined to contribute to the inner cell mass (White et al., 2016). We propose that SOX2 is associated with future pluripotent state but does not alone contribute to all aspects of pluripotency, such as inside positioning. It is therefore still unclear why it is important to establish the inside cell-specific SOX2 expression during embryogenesis. Identification pathways that function downstream of YAP1/WWTR1 and in parallel to SOX2 to promote formation of pluripotent cells will provide meaningful insights into the natural origins of mammalian pluripotent stem cell progenitors.

## Methods

### Mouse strains and genotyping

All animal research was conducted in accordance with the guidelines of the Michigan State University Institutional Animal Care and Use Committee. Wild type embryos were derived from CD-1 mice (Charles River). The following alleles or transgenes were used in this study, and maintained in a CD-1 background: *Sox2^tm1.1Lan^* (Smith et al., 2009), *Yap^tm1.1Eno^* (Xin et al., 2011), *Wwtr1^tm1.1Eno^* (Xin et al., 2013), *Tg(Zp3-cre)93Knw* (de Vries et al., 2000). Null alleles were generated by breeding mice carrying floxed alleles and mice carrying ubiquitously expressed *Cre*, *129-Alpl^tm(cre)Nagy^* (Lomelí et al., 2000).

### Embryo collection and culture

Mice were maintained on a 12-hour light/dark cycle. Embryos were collected by flushing the oviduct or uterus with M2 medium (Millipore). For embryo culture, KSOM medium (Millipore) was equilibrated overnight prior to embryo collection. Y-27632 (Millipore) was included in embryo culture medium at a concentration of 80 μM with 0.4% DMSO, or 0.4% DMSO as control, where indicated. Embryos were cultured at 37°C in a 5% CO_2_ incubator under light mineral oil.

### Embryo microinjection

LATS2 and YAPS112A mRNA was synthesized from cDNAs cloned into the pcDNA3.1-poly(A)83 plasmid (Yamagata et al., 2005) using the mMESSAGE mMACHINE T7 transcription kit (Invitrogen). *EGFP* or *nls-GFP* mRNA were synthesized from *EGFP* cloned into the pCS2 plasmid or the *nls-GFP* plasmid (Ariotti et al., 2015) using the mMESSAGE mMACHINE SP6 transcription kit (Invitrogen). mRNAs were cleaned and concentrated prior to injection using the MEGAclear Transcription Clean-Up Kit (Invitrogen). *Lats2*, *Lats2^KD^* and *YAP^CA^* mRNAs were injected into one blastomere of two-cell stage embryos at a concentration of 500 ng/μl, mixed with 350 ng/μl *EGFP* or *nls-GFP* mRNA diluted in 10 mM Tris-HCl (pH 7.4), 0.1 mM EDTA.

### Immunofluorescence and Confocal Microscopy

Embryos were fixed with 4% formaldehyde (Polysciences) for 10 minutes, permeabilize with 0.5% Triton X-100 (Sigma Aldrich) for 30 minutes, and then blocked with blocking solution (10% Fetal Bovine Serum (Hyclone), 0.2% Triton X-100) for 1 hour at room temperature, or overnight at 4°C. Primary Antibodies used were: mouse anti-CDX2 (Biogenex, CDX2-88), goat anti-SOX2 (Neuromics, GT15098), rabbit anti-PARD6B (Santa Cruz, sc-67393), rabbit anti-PARD6B (Novus Biologicals, NBP1-87337), mouse anti-PKCζ (Santa Cruz Biotechnology, sc-17781), rat anti-CDH1 (Sigma Aldrich, U3254), mouse anti-YAP (Santa Cruz Biotechnology, sc101199), rabbit anti phospho-YAP (Cell Signaling Technologies, 4911), chicken anti-GFP (Aves, GFP-1020). Stains used were: Phallodin-633 (Invitrogen), DRAQ5 (Cell Signaling Technologies) and DAPI (Sigma Aldrich). Secondary antibodies conjugated to DyLight 488, Cy3 or Alexa Flour 647 fluorophores were obtained from Jackson ImmunoResearch. Embryos were imaged using an Olympus FluoView FV1000 Confocal Laser Scanning Microscope system with 20x UPlanFLN objective (0.5 NA) and 5x digital zoom. For each embryo, z-stacks were collected, with 5 μm intervals between optical sections. All embryos were imaged prior to knowledge of their genotypes.

### Embryo Analysis

For each embryo, z-stacks were analyzed using Photoshop or Fiji, which enabled the virtual labeling, based on DNA stain, of all individual cell nuclei. Using this label to identify individual cells, each cell in each embryo was then assigned to relevant phenotypic categories, without knowledge of embryo genotype. Phenotypic categories included marker expression (e.g., SOX2 or CDX2 positive or negative), protein localization (e.g., aPKC or CDH1 apical, basal, absent, or unlocalized), and cell position, where cells making contact with the external environment were considered ‘outside’ and cells surrounded by other cells were considered ‘inside’ cells.

### TUNEL Assay

Embryos were fixed, permeabilized, and blocked as described for immunofluorescence. Zonae pellucida were removed using Tyrode’s Acid treatment prior to performing the TUNEL assay (*In Situ* Cell Death Detection Kit, Fluorescein, Millipore-Sigma). Embryos were incubated in 200 μl of a 1:10 dilution of enzyme in label solution for 2 hours at 37 °C. Embryos were then washed twice with blocking solution for 10 minutes each, and then mounted in a 1 to 400 dilution of DRAQ5 in blocking solution to stain DNA.

### Embryo Genotyping

To determine embryo genotypes, embryos were collected after imaging and genomic DNA extracted using the Extract-N-Amp kit (Sigma) in a final volume of 10 μl. Genomic extracts (1-2 μl) were then subjected to PCR using allele-specific primers (Table S3).

## Acknowledgements

We are grateful to Dr. Hiroshi Sasaki for providing expression constructs, to Dr. Randy L. Johnson for providing mice carrying conditional alleles of *Yap1* and *Wwtr1,* and to Dr. Jason Knott for embryo microinjection training. We also thank Dr. Ripla Arora, Dr. Julia Ganz, and members of the Ralston Lab for comments. This work was supported by NIH R01 GM104009 and the Eunice Kennedy Shriver National Institute of Child Health & Human Development of the National Institutes of Health under Award Number T32HD087166. The content is solely the responsibility of the authors and does not necessarily represent the official views of the National Institutes of Health. We thank anonymous reviewers for insightful questions and suggestions.

---

**Figure S1.**
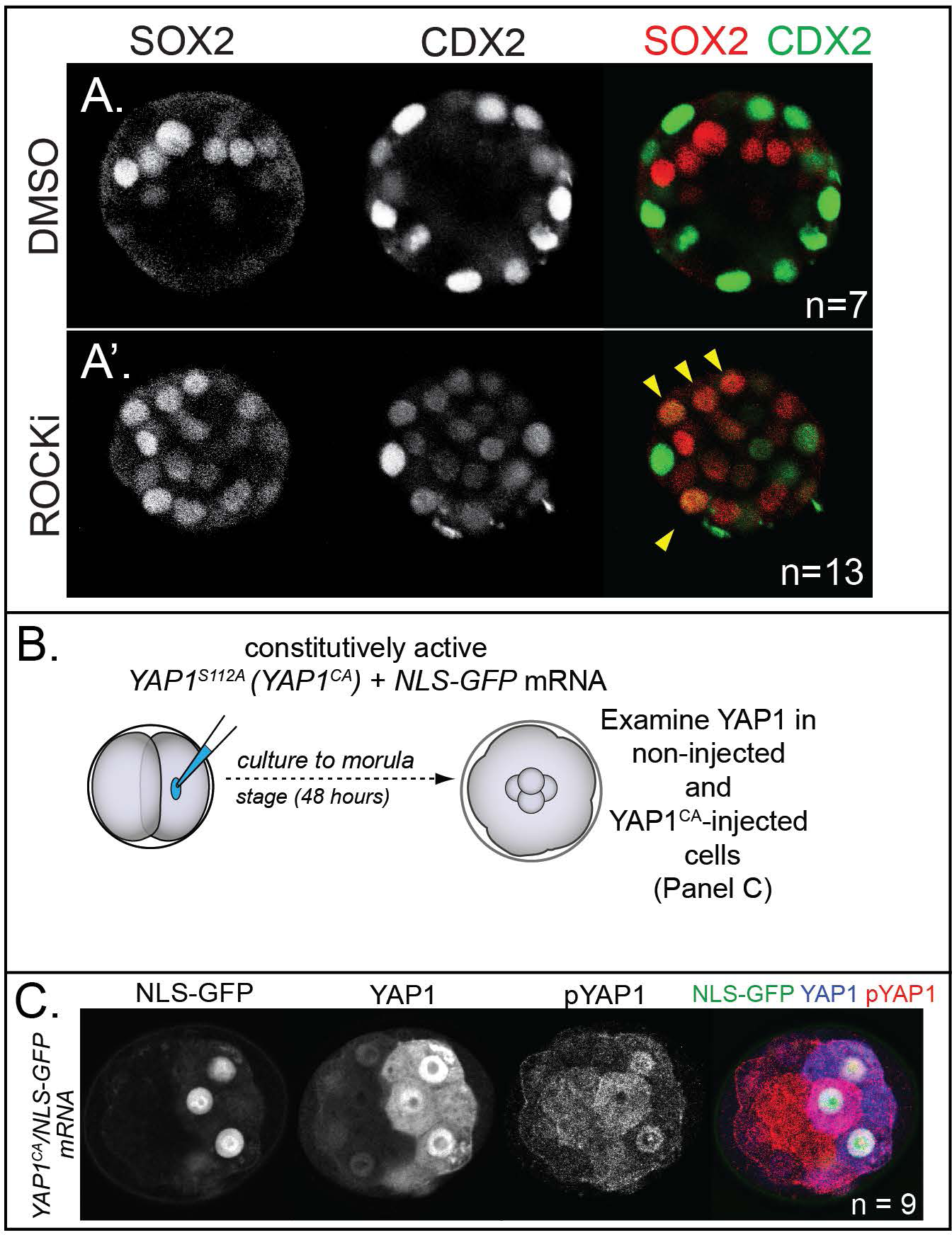
Effect of ROCK1/2 inhibition on *Cdx2* expression and effect *Yap1^CA^* overexpression on YAP1 localization and phosphorylation. A) Confocal images of CDX2 and SOX2 in control and embryos treated with ROCKinhibitor for 24 hours starting at E2.5. In control embryos, CDX2 is specific to outside cells and SOX2 is specific to inside cells (n = embryos). A’) Treatment with ROCKi leads to ectopic SOX2 in outside cells which is often co expressed with CDX2 (arrowheads, n = embryos). B) *YAP1^CA^* was injected into one of two blastomeres at the 2-cell stage and evaluated 48 hours later. C) In non-injected cells, YAP1 is exclusively nuclear in outside cells while pYAP1 is exclusively cytoplasmic in inside cells. By contrast, YAP1 is detected in the nucleus of *YAP1^CA^*-injected cells, regardless of their position, demonstrating that YAP1^CA^ is constitutively nuclear. Additionally, analysis of pYAP1 in *YAP1^CA^*-injected cells shows that *YAP1^cA^* can still be phosphorylated on non-mutated residues, but this is not sufficient to alter YAP1 nuclear localization (n = embryos).

**Figure S2.**
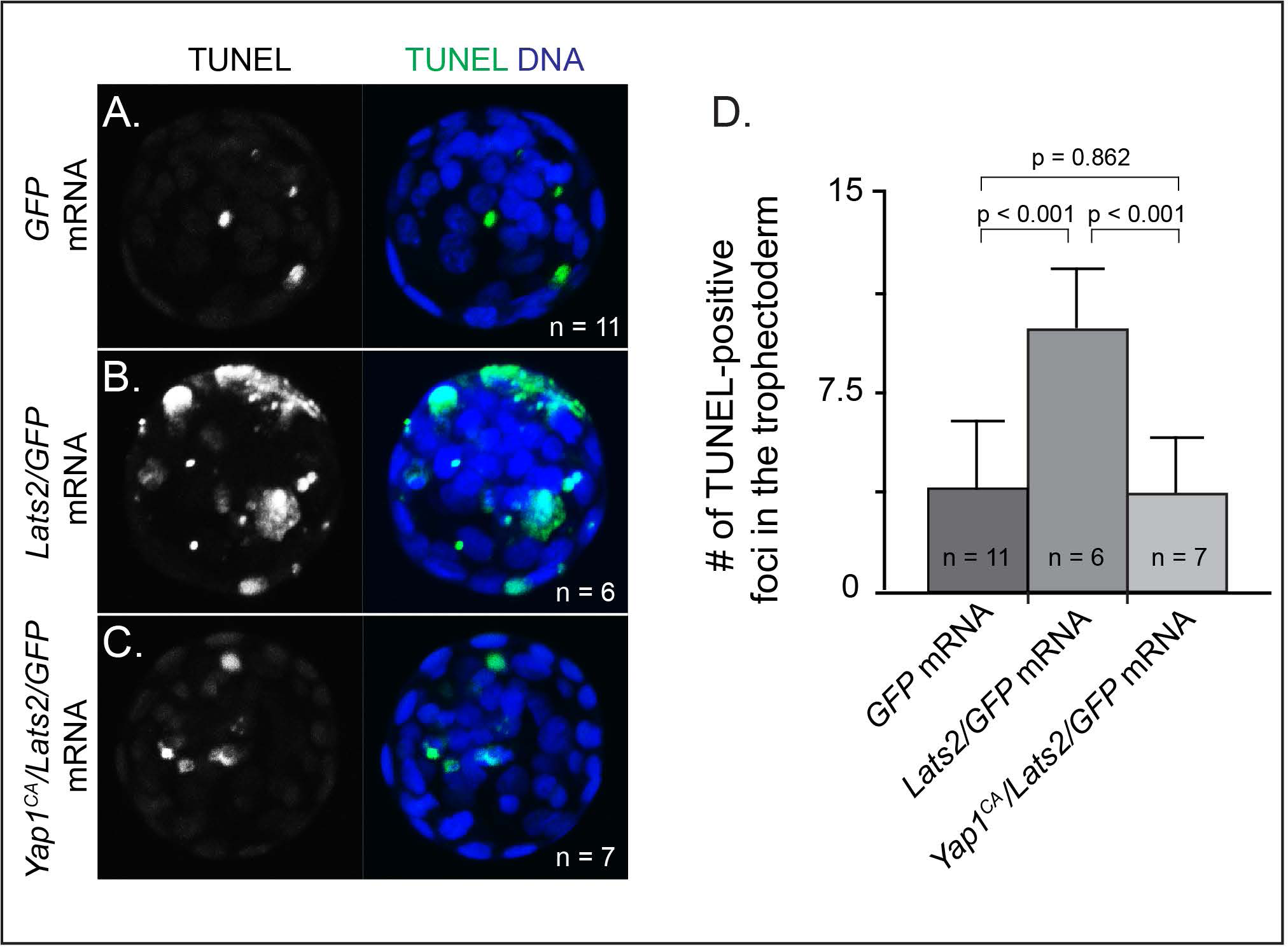
*Lats2*-overexpressing cells die on the surface of the embryo. A) Merge of all confocal sections from TUNEL assay performed on an embryo injected with *GFP* mRNA into one blastomere at the two-cell stage and then cultured until the blastocyst stage. Note that GFP fluorescence does not survive the TUNEL assay (n = embryos). B) Embryo injected in one of two cells with *Lats2* and *GFP* mRNA and then cultured until the blastocyst stage shows elevated TUNEL-positive foci in the outside cells (n = embryos). C) Embryo injected in one of two cells with *Yap1^CA^*, *Lats2* and *GFP* mRNA and then cultured until the blastocyst stage shows reduced TUNEL-positive foci in the outside cells (n = embryos). D) Quantification, across indicated sample sizes, of the average number of TUNEL-positive foci per embryo (t = student’s t-test, n = embryos).

**Figure S3.**
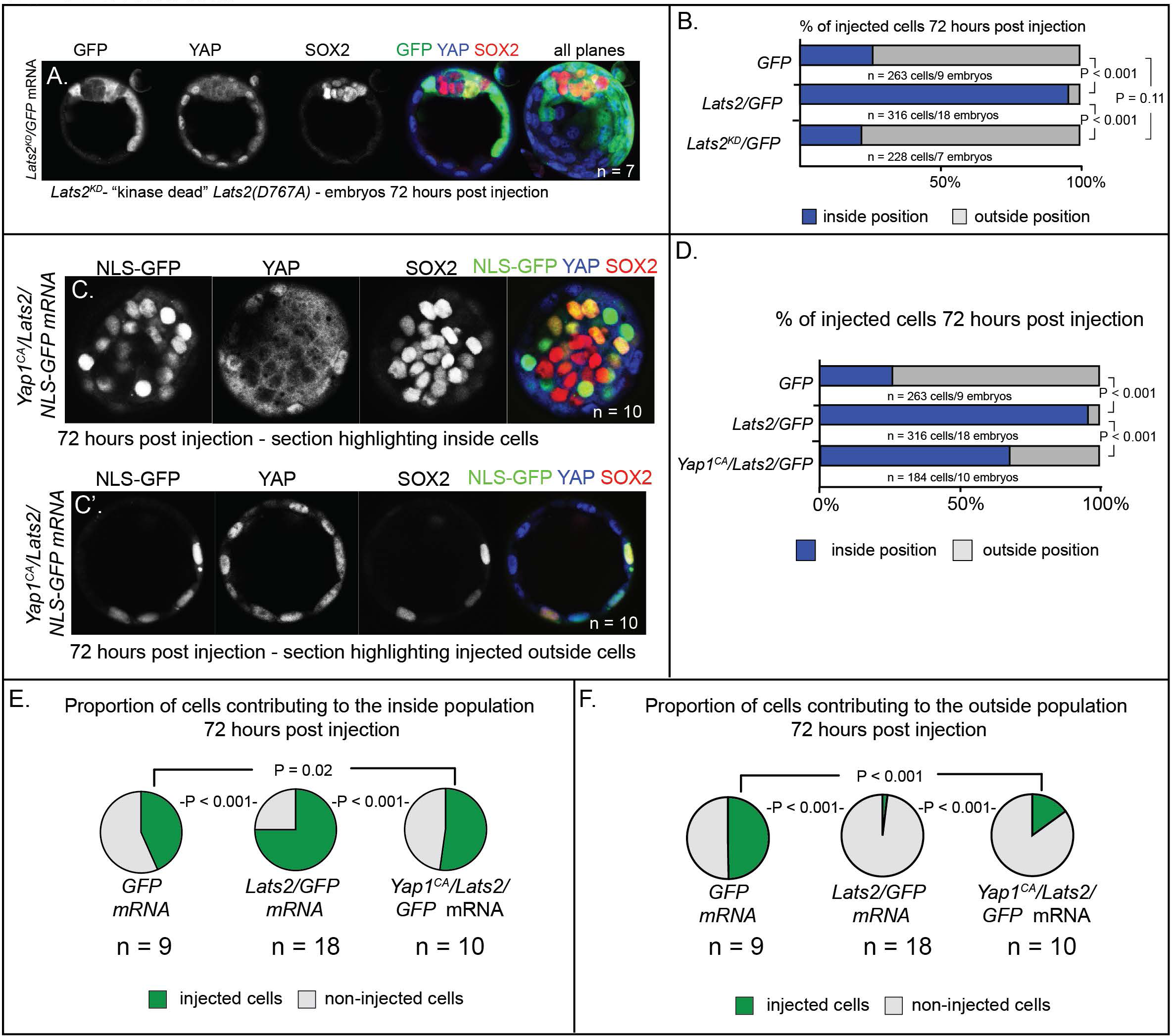
LATS2 drives cells to an inside position by inhibiting YAP1 activity. A) *Lats2^KD^/GFP*-injected cells contribute to trophectoderm and inner cell mass and do not alter expression of SOX2 (n = embryos). B) Contribution of injected cells to inside and outside embryo compartments. Unlike *Lats2*-injected cells, *Lats2^KD^-*injected cells do not show an increased tendency to localize to the inside of the embryo (P, chi-squared test). C-C’) Cooverexpression of *Yap1^CA^* and *Lats2* partially rescues the ability of Lats2-overexpressing cells to contribute to trophectoderm and to repress *Sox2.* Panel (C) shows a confocal section that includes the inside cell population of a *Yap1^CA^/Lats2* injected embryo, showing inhibition by *Yap1^CA^* on *Sox2* expression in some Lats2-overexpressing inside cells. Panel (C’) shows a confocal section of the same embryo and highlights the contribution of cells cooverexpressing and highlights the contribution of cells overexpressing *Yap1^CA^* and *Lats2* to the trophectoderm (n = embryos). D) Contribution of injected cells to inside and outside embryo compartments. *Yap1^CA^-* overexpression partially reverses the effect of *Lats2* overexpression on cellular localization to outside and inside regions of the embryo (P, chi-squared test). E) Proportion of non-injected cells and injected cells contributing to the inside population in embryos injected with the indicated mRNAs. Yap1^CA^-cooverexpression reduces the proportion of Lats2-overexpressing cells observed in the inside population (P, chi-squared test, n = embryos). F) Proportion of non-injected cells and injected cells contributing to the outside population in embryos injected with the indicated mRNAs. *Yap1^CA^*-cooverexpression increases the proportion of *Lats2*-overexpressing cells observed in the outside population (P, chi-squared test, n = embryos).

**Figure S4.**
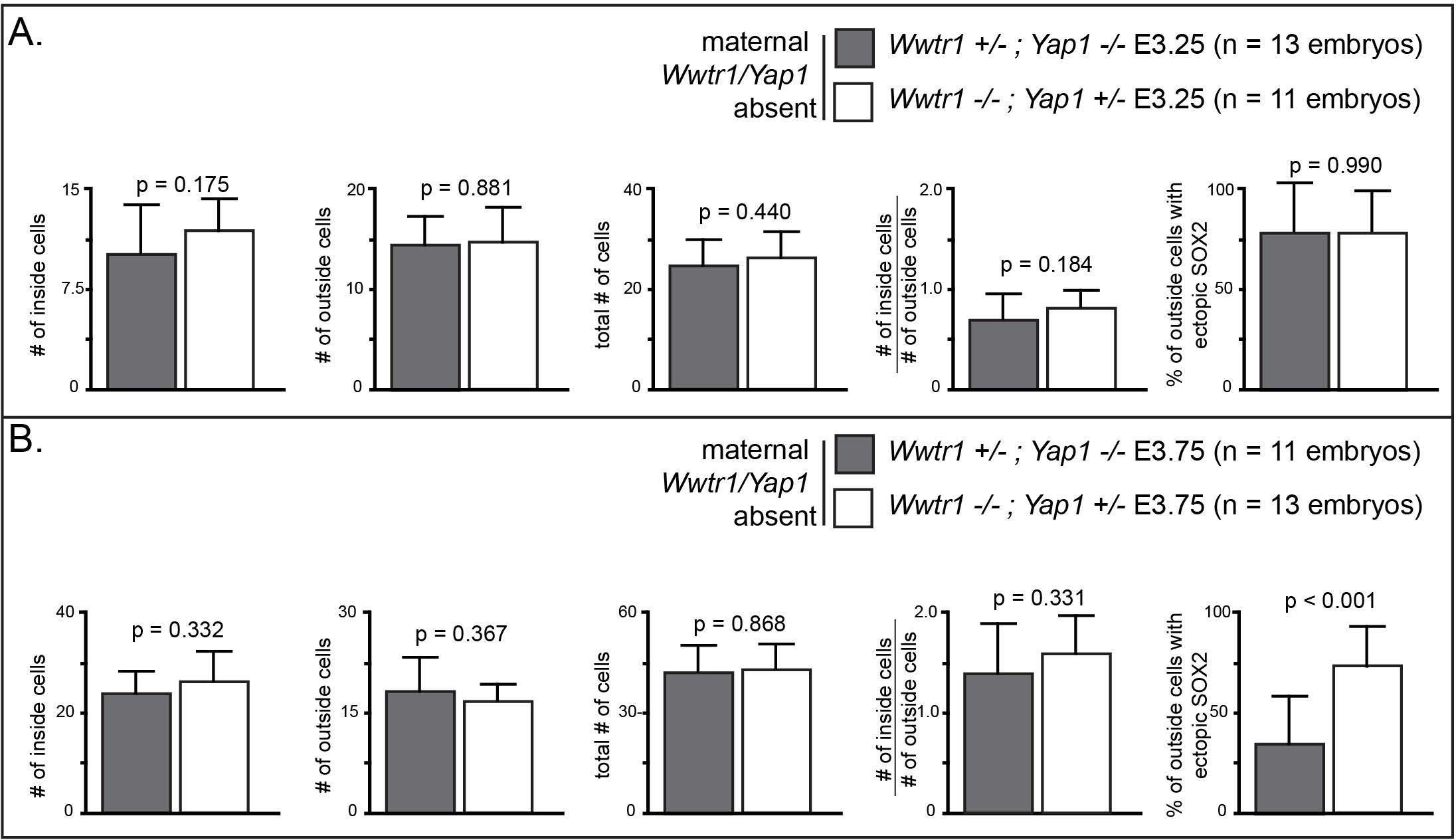
*Wwtr1* and *Yap1* are functionally equivalent during cell positioning, but not repression of *Sox2* in outside cells at E3.75. A) Average numbers per embryo of each stated category in embryos of indicated genotypes at E3.25. No differences were detected between the two genotypes at this stage (p, student’s t-test; n = embryos). B) Average numbers per embryo of each stated category in embryos of indicated genotypes at E3.75. The only significant difference observed was in the degree of ectopic SOX2 detected in outside cells, a phenotype that was more severe in embryos lacking *Wwtr1* (p, Student’s t-test; n = embryos).

**Figure S6.**
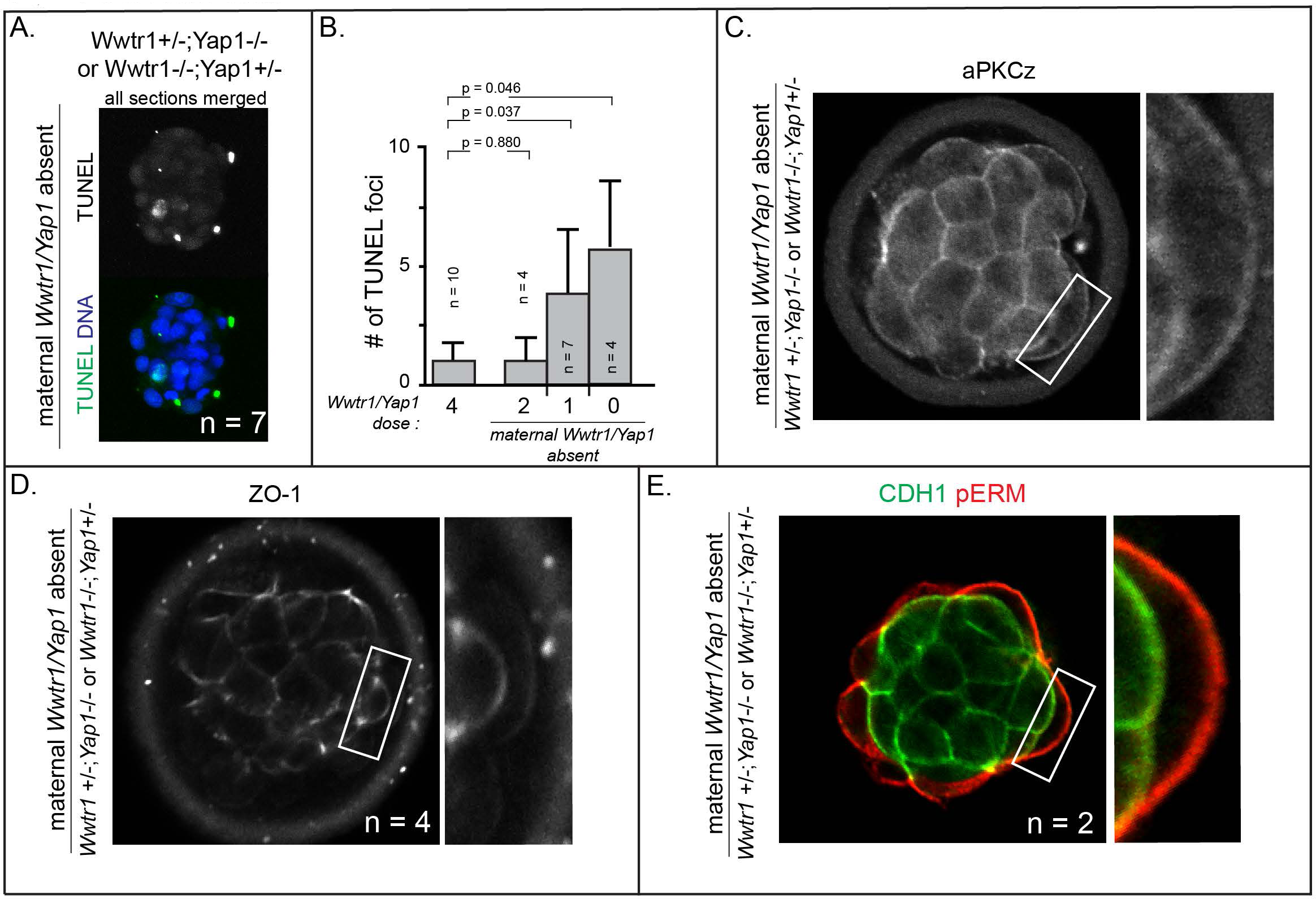
Increased cell death and epithelialization defects in embryos lacking maternal *Wwtr1* and *Yap1* with a single wild type allele of *Wwtr1* or *Yap1.* A) TUNEL staining in embryos lacking maternal *Wwtr1* and *Yap1* with a single wild type allele of *Wwtr1* or *Yap1.* Max projection of all confocal sections taken from a single embryo is shown (n = embryos) B) Quantification of the average number of TUNEL foci per embryo in embryos lacking with decreasing doses of *Wwtr1* and *Yap1* (n = embryos) C) aPKCz staining in embryos lacking maternal *Wwtr1* and *Yap1* with a single wild type allele of *Wwtr1* or *Yap1.* aPKC is not localized to the apical membrane of embryos lacking maternal *Wwtr1* and *Yap1* with a single wild type allele of *Wwtr1* or *Yap1* (n = embryos). D) ZO-1 staining in embryos lacking maternal *Wwtr1* and *Yap1* with a single wild type allele of *Wwtr1* or *Yap1.* aPKC is not localized to the apical membrane of embryos lacking maternal *Wwtr1* and *Yap1* with a single wild type allele of *Wwtr1* or *Yap1* (n = embryos). E) pERM and CDH1 staining in embryos lacking maternal *Wwtr1* and *Yap1* with a single wild type allele of *Wwtr1* or *Yap1.* pERM is correctly localized to apical membranes and CDH1 correctly localized to basolateral membranes in all embryos (n = embryos).

**Table S1.**
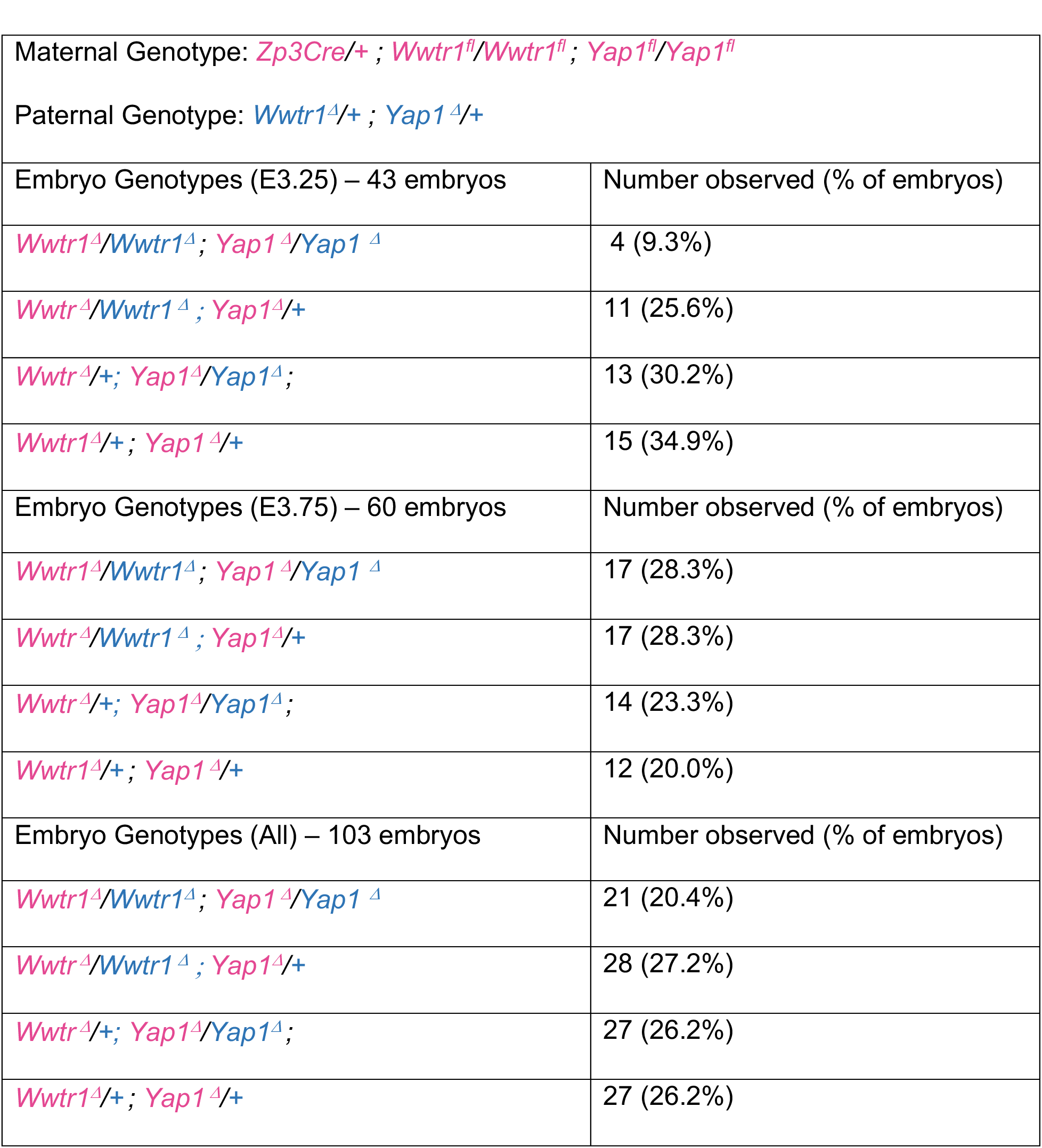
Summary of embryos recovered from *Wwtr1;Yap1* germline null females

**Table S2:** Mean and standard deviation of cell counts for every experimental treatment

|  |  |  |  |  |
| --- | --- | --- | --- | --- |
| Figure 1 and S1 | INSIDE | OUTSIDE | TOTAL | INSIDE/OUTSIDE ratio |
| Control (n = 17) | 14.6 +/- 4.2 | 23.0 +/- 7.1 | 37.7 +/- 10.4 | 0.66 +/- 0.14 |
| ROCK-inhibitor treated (n = 23) | 10.8 +/- 4.7 | 24.6 +/- 5.3 | 35.3 +/- 8.8 | 0.44 +/- 0.14 |
| Yap1CA/GFP (n = 10) | 18.4 +/- 7.3 | 36.6 +/- 8.9 | 55.0 +/- 10.3 | 0.54 +/- 0.24 |
| Figures 2 and S3 | INSIDE | OUTSIDE | TOTAL | INSIDE/OUTSIDE ratio |
| GFP Only (n = 9) | 15.6 +/- 3.1 | 44.4 +/- 12.8 | 60.1 +/- 12.9 | 0.39 +/- 0.17 |
| Lats2/GFP (n = 18) | 22.9 +/- 7.3 | 32.7 +/- 11.0 | 55.6 +/- 12.3 | 0.79 +/- 0.39 |
| Lats2KD/GFP (n = 7) | 17.4 +/- 4.0 | 51.4 +/- 8.8 | 68.9 +/- 12.2 | 0.34 +/- 0.05 |
| Yap1CA/Lats2/GFP (n = 9) | 25.4 +/- 7.0 | 39.1 +/- 10.7 | 64.6 +/- 9.5 | 0.71 +/- 0.29 |
| Figure 3 | INSIDE | OUTSIDE | TOTAL | INSIDE/OUTSIDE ratio |
| Sox2 m null Lats2/GFP (n = 5) | 23 +/- 4.6 | 32.8 +/- 8.8 | 55.8 +/- 12.4 | 0.72 +/- 0.15 |
| Sox2 m/z null Lats2/GFP (n = 5) | 19.8 +/- 3.3 | 37.4 +/- 14.6 | 57.2 +/- 13.6 | 0.58 +/- 0.19 |
| Figure 4 | INSIDE | OUTSIDE | TOTAL | INSIDE/OUTSIDE ratio |
| GFP Only (n = 18) | 5.8 +/- 4.9 | 14.8 +/- 3.0 | 20.8 +/- 6.4 | 0.38 +/- 0.31 |
| Lats2/GFP (n = 58) | 4.2 +/- 3.8 | 13.4 +/- 3.5 | 17.2 +/- 6.3 | 0.31 +/- 0.25 |
| Figure 5 | INSIDE | OUTSIDE | TOTAL | INSIDE/OUTSIDE ratio |
| wild type E3.25 (n = 3) | 10.6 +/- 1.5 | 16.6 +/- 2.1 | 27.7 +/- 1.2 | 0.65 +/- 0.17 |
| Wwtr1 +/-;Yap1 +/- E3.25 (n = 15) | 9.1 +/- 3.0 | 13.6 +/- 3.6 | 22.6 +/- 5.4 | 0.69 +/- 0.23 |
| Wwtr1 +/-;Yap1 -/- or Wwtr1 -/-;Yap1 +/- E3.25 (n = 24) | 11.0 +/- 3.2 | 14.7 +/- 3.1 | 25.6 +/- 5.2 | 0.77 +/- 0.22 |
| Wwtr1 -/-;Yap1 -/- E3.25 (n = 4) | 12.0 +/- 4.2 | 13.5 +/- 4.4 | 25.4 +/- 4.0 | 0.98 +/- 0.51 |
| Figures 6 and S6 | INSIDE | OUTSIDE | TOTAL | INSIDE/OUTSIDE ratio |
| wild type E3.75 (n = 8) | 16.5 +/- 1.8 | 41.8 +/- 6.8 | 58.3 +/- 7.0 | 0.29 +/- 0.04 |
| Wwtr1 +/-;Yap1 +/- E3.75 (n = 8) | 17.6 +/- 5.0 | 31.4 +/- 13.6 | 50.3 +/- 13.9 | 0.36 +/- 0.11 |
| Wwtr1 +/-;Yap1 -/- or Wwtr1 -/-;Yap1 +/- E3.75 (n = 24) | 25.4 +/- 5.3 | 17.5 +/- 4.0 | 42.9 +/- 7.3 | 0.59 +/- 0.07 |
| Wwtr1 -/-;Yap1 -/- E3.75 (n = 13) | 20.8 +/- 2.96 | 15.2 +/- 3.6 | 36.0 +/- 6.2 | 0.58 +/- 0.04 |

**Table S3.**
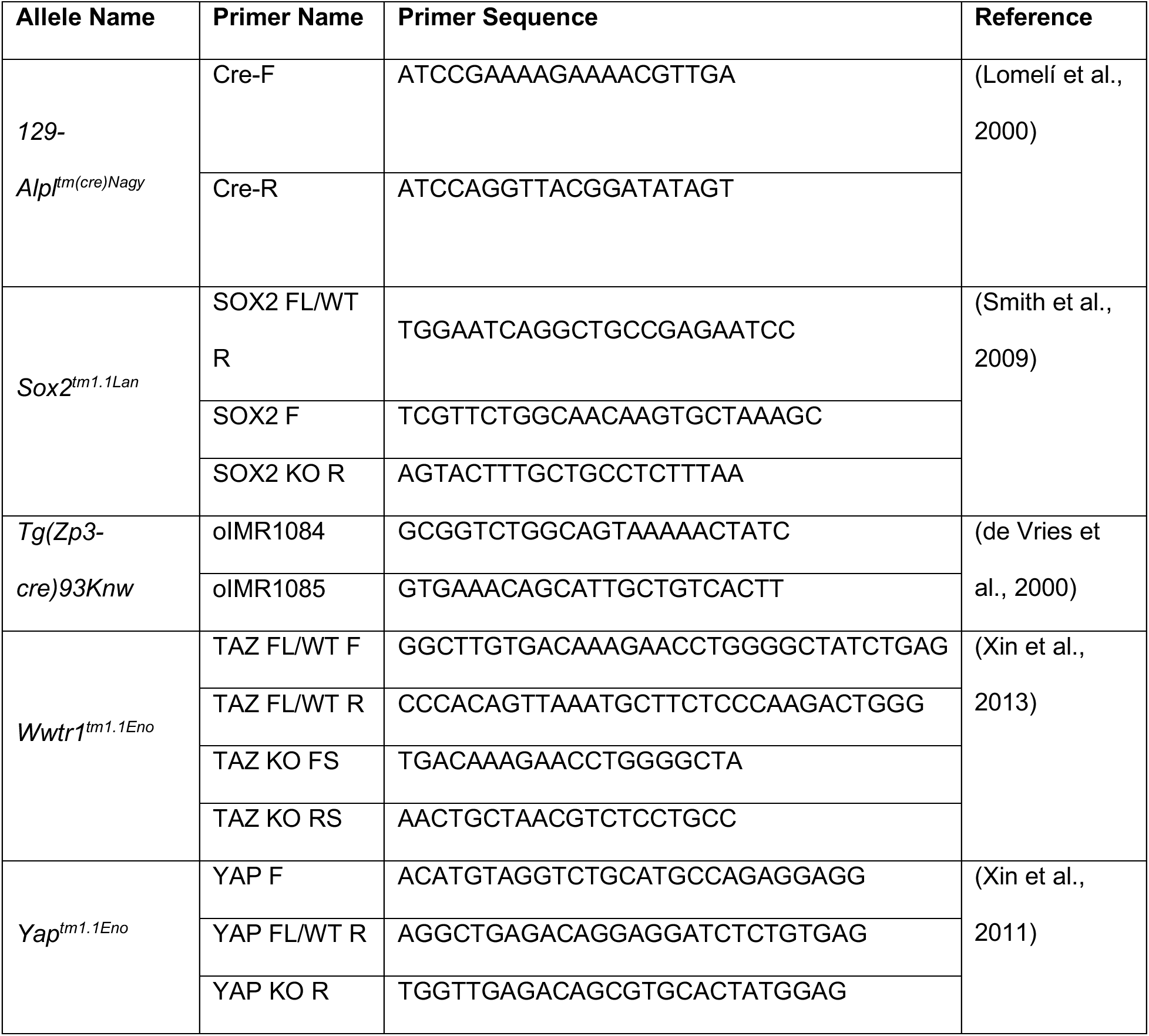
Allele-specific primers used for determining embryo and mouse genotypes

